# A Complex-valued Oscillatory Neural Network for Storage and Retrieval of Multichannel Electroencephalogram Signals

**DOI:** 10.1101/2020.03.28.013292

**Authors:** Dipayan Biswas, P. Sooryakiran, V. Srinivasa Chakravarthy

## Abstract

Recurrent neural networks with associative memory properties are typically based on fixed-point dynamics, which is fundamentally distinct from the oscillatory dynamics of the brain. There have been proposals for oscillatory associative memories, but here too, in the majority of cases, only binary patterns are stored as oscillatory states in the network. Oscillatory neural network models typically operate at a single/common frequency. At multiple frequencies, even a pair of oscillators with real coupling exhibits rich dynamics of Arnold tongues, not easily harnessed to achieve reliable memory storage and retrieval. Since real brain dynamics comprises of a wide range of spectral components, there is a need for oscillatory neural network models that operate at multiple frequencies. We propose an oscillatory neural network that can model multiple time series simultaneously by performing a Fourier-like decomposition of the signals. We show that these enhanced properties of a network of Hopf oscillators become possible by operating in the complex-variable domain. In this model, the single neural oscillator is modeled as a Hopf oscillator, with adaptive frequency and dynamics described over the complex domain. We propose a novel form of coupling, dubbed “power coupling,” between complex Hopf oscillators. With power coupling, expressed naturally only in the complex-variable domain, it is possible to achieve stable (normalized) phase relationships in a network of multifrequency oscillators. Network connections are trained either by Hebb-like learning or by delta rule, adapted to the complex domain. The network is capable of modeling N-channel Electroencephalogram time series with high accuracy and shows the potential as an effective model of large-scale brain dynamics.

## 1. Introduction

Currently, there are two prominent approaches to characterizing the neural code: the instantaneous rate or frequency at which the neuron fires action potential (“the spike frequency code” or the “rate code”) and the time of the occurrence of action potential (“spike time code”). The former assumes the information is processed/encoded over a larger time scale conveyed by average number of spikes fired in a given duration forms the basis of a large class of neural networks called the rate coded neural networks (Ruck et al., 1990; Lawrence et al., 1997; Lippmann, 1990). Whereas the latter believe that the precise timing of the action potential fired by a neuron encodes the ongoing activity, giving rise to a broad class of spiking neural network (Maass, 1997; Izhikevich, 2003; Izhikevich, 2004; Ghosh-dastidar and Lichtenstein, 2009). These two classes of neural networks are capable of universal function approximation and well explored (Maass and Maass, 1997; Auer et al., 2008). They have been reported to solve a wide range of information processing problems like vector space transformation, dimensionality reduction, sequence processing, memory storage as an attractor and autoencoding (Schmidhuber, 2015; Kohonen, 1998; Frasconi et al., 1995; Hopfield, 1982; Trappenberg, 2003).

The third class of neural code, dubbed the oscillation coding (Fujii et al 1996) also has been around for more than two decades. However, it has not been adopted in modeling literature as extensively as the first two types of neural code. Oscillation coding is based on the idea that not single-neuron activity, but the collective activity of a neural ensemble is the true building block of the brain’s activity. Unlike the rate code, which is often represented as a binary signal, or the spike time code, which is often described as a train of delta functions, oscillation code is a smooth signal that is said to be composed of distinct frequency bands. Oscillation coding also offers an opportunity to represent important temporal phenomena like temporal binding via transient synchrony (Wang and Terman, 1995; Wang and Terman, 1997; Shi et al., 2008) or rhythmic behaviors such as locomotion (ERMENTROUT and KOPELL, 1989). There have been comprehensive and valorous attempts to describe all brain function in terms of the oscillatory activity of neural ensembles (Buzsáki et al., 2012; Draguhn, 2004). However, since existing literature does not commit to the exact size of a ‘neural ensemble,’ activity measured at different scales goes by different names, including local field potentials (LFPs), electrocorticograms (ECoGs), electroencephalograms (EEGs), etc.

Since oscillations are ubiquitous in the brain, one would naturally expect nonlinear oscillator models to be used extensively to describe brain function. However, models of brain function often use sigmoidal neurons (to represent the rate code), spiking neuron models (to represent the spike time code), or more detailed conductance-based or biophysical models. In the field of neural signal processing, for example, a large body of literature depicting various deep neural network architectures such as recurrent neural network, convolutional neural network and deep belief network have been used for EEG signal processing and classification at various scales which found application in the following areas: brain-computer interface, epilepsy, cognitive and effective monitoring, etc. (Craik and He, 2019; Roy et al., 2019). Nonlinear oscillators are used exclusively for describing oscillatory phenomena like rhythmic movements or feature binding by synchronization. In an attempt to correct this bias, it would be a worthwhile modeling exercise to construct general networks of nonlinear neural oscillators that can be used to describe a wide range of brain functions.

In this paper, we describe a general trainable network of neural oscillators. For the oscillator model, we choose the simple Hopf oscillator. One reason behind the choice of the Hopf oscillator is the elegant form its description assumes in the complex variable domain. We show that it is important to operate in the complex domain in order to define some of the novel modeling features we introduce in this paper. Particularly, we introduce the concept of ‘power coupling’ by which it is possible to achieve robust phase relationships among oscillators with very different intrinsic frequencies. The proposed network of oscillators can be trained to model multi-channel EEG, thereby demonstrating the potential to evolve into a large-scale brain model.

The outline of the paper is as follows. Section 2 begins with the definition of Hopf oscillator in the complex domain and show how to adapt the intrinsic frequency of the oscillator to that of a sinusoidal forcing signal. Section 2.4 describes the dynamics of a pair of coupled Hopf oscillators and shows the advantages in coupling with a complex coupling constant. Section 2.6 highlights the difficulties in coupling a pair of Hopf oscillators with very different intrinsic frequencies, and Section 2.7 shows how the difficulties can be overcome by adopting “power coupling.” Gathering all the modeling elements developed so far, Section 2.12 describes a network of oscillators that can model multi-channel EEG. A discussion of the work was presented in the last section.

## 2. A network of complex-valued neural oscillators

### 2.1 Hopf oscillator in the complex form

The canonical model of Hopf oscillator without any external input is described as follows in complex state variable representation 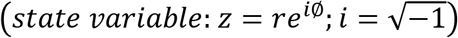, Cartesian coordinate representation (*state variables: x,y*) and polar coordinate representation (*state variables: r*, 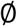) respectively:

Complex state variable representation:

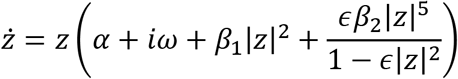

Cartesian coordinate representation:

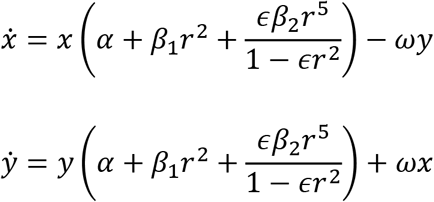

Polar coordinate representation:

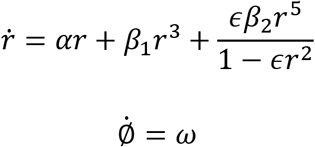

Depending on the parameter values (*α,β*_1_,*β*_2_), the autonomous behavior of the oscillator principally falls into four regimes: critical Hopf regime (*α = 0,β*_1_ < 0, *β*_2_ = 0), supercritical Hopf regime (*α* > 0, *β*_1_ < 0, *β*_2_ = 0), supercritical double limit cycle regime (*α* <0, *β*_1_> 0, *β*_2_ < 0, *local maxima* > 0) and subcritical double limit cycle regime (*α* < 0, *β*_1_ > 0, *β*_2_ < 0, *local maxima* < 0) (Kim and Large, 2015). The critical Hopf regime has a stable fixed point at the origin and has the ability to show a stable resonating response to the complex sinusoidal input (*Fe^iω_0_t^*). The supercritical Hopf regime has an unstable fixed point at the origin and a stable limit cycle at 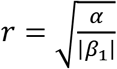. In the supercritical double limit cycle regime, the system exhibits two limit cycles, one of which is stable while the other being unstable. In the subcritical Hopf regime, the system has one stable fixed point at the origin. However, it has the ability to show stable oscillation under the influence of complex sinusoidal input whose frequency is not too different from that of the oscillator (*ω* – *ω*_0_) < *σ*. Throughout the rest of the paper, we will be using supercritical Hopf regime, which can be defined as follows:

Complex state variable representation:

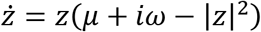

Cartesian coordinate representation:

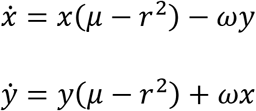

Polar coordinate representation:

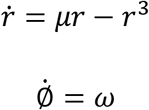

We now present a series of results related to a single oscillator, a coupled pair, and a network of Hopf oscillators in the supercritical regime defined above.

### 2.2 Single oscillator with adaptive frequency

It has been previously shown that when a Hopf oscillator is influenced by a real sinusoidal input signal, it can adapt its natural frequency to the frequency of the input signal if it follows the following dynamics (Righetti et al., 2005).

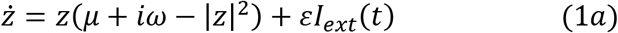

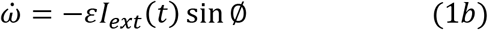

Given, 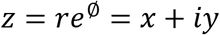. It can be recognized that the Hopf oscillator is perturbed by a real input signal *I_ext_*(*t*) with coupling strength *ε*. At steady-state *r* reaches 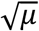 for *ε* = 0. If *I_ext_*(*t*) = *A* sin(*ω*_0_*t* + *φ*), the natural frequency, *ω*, adapts to the frequency *ω*_0_.

#### Adaptive Hopf oscillator with complex input or complex adaptive Hopf oscillator

When a Hopf oscillator is influenced by a complex sinusoidal input signal, the natural frequency of the oscillator can adapt to the frequency of input if the natural frequency of the oscillator is updated according to the following equation.

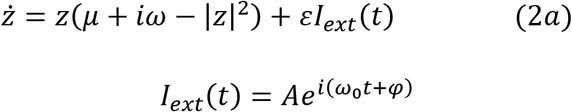

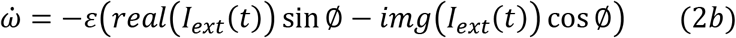

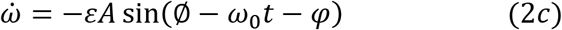

In this scenario, *I_ext_*(*t*) is a complex sinusoidal signal. It is straight forward to derive the learning rule for the natural frequency of the oscillator if equation-2a is represented in the Cartesian and polar coordinate forms, respectively, as follows.

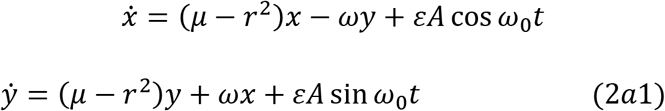

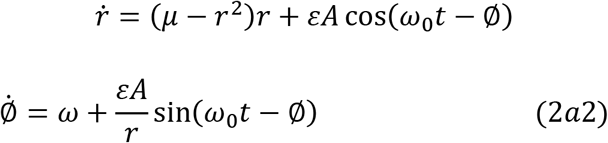

In the phase plane representation, it can be observed from equation-2a2 that the influence caused by the input perturbation on the oscillator phase is 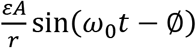. Whereas from the Cartesian coordinate system representation (equation-2a1) it can be observed that the overall influence of the external perturbation on the phase point along the tangential axis of the limit cycle 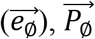 is (to understand the vector notations refer to fig-2):

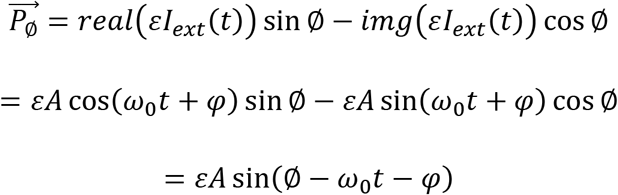

**Fig-1:**
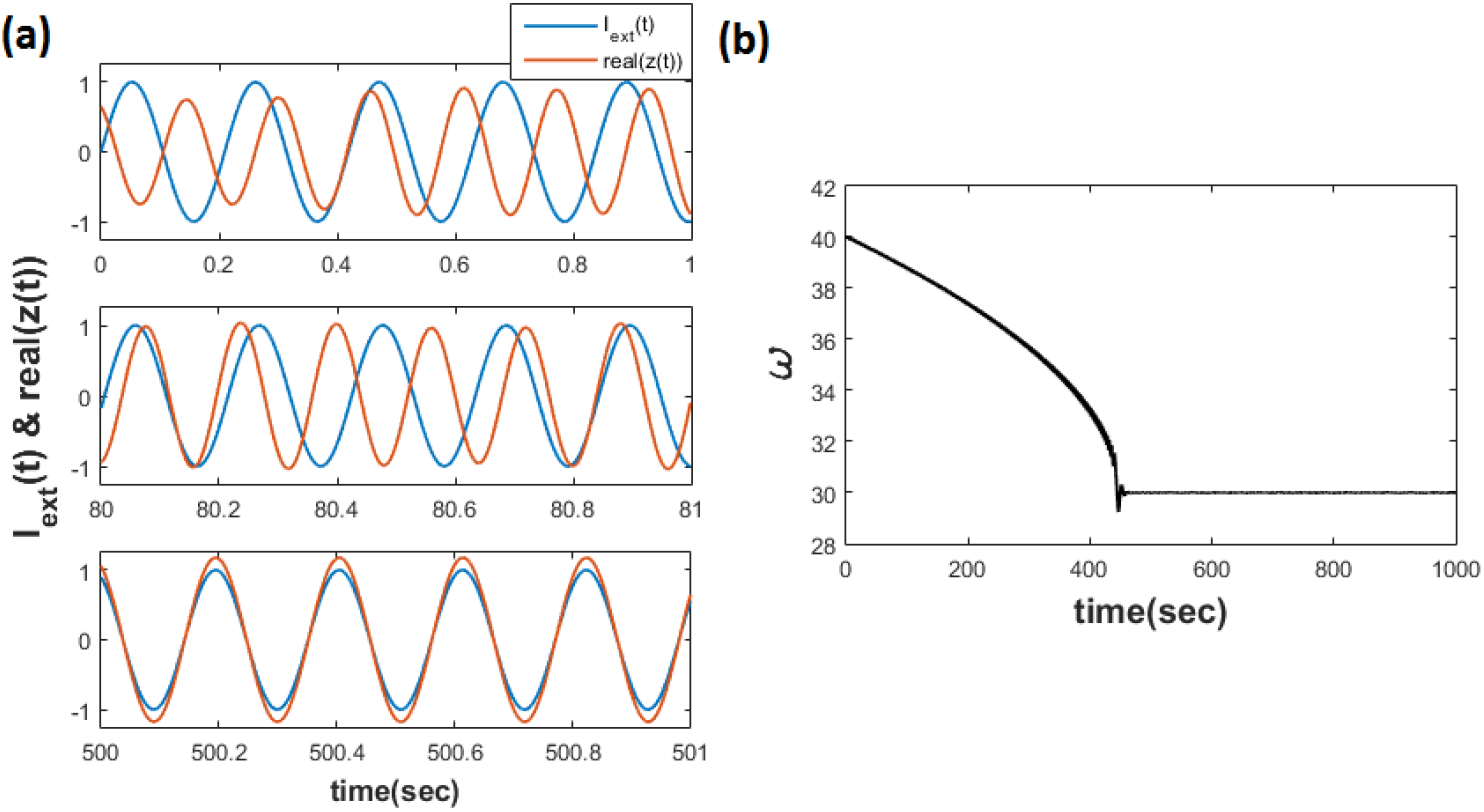
Equation-1 is simulated for *μ* = 1, *ω*(0) = 40, *ω*_0_ = 30, *ε* = 0.9, *A* = 1, *φ* = 0 for 1000 secs. (a) Depicts the variation of *x*(*t*) w.r.t *I_ext_*(*t*) at various time instants. (b) It can be observed that the natural frequency of the Hopf oscillator adapts to the frequency of the input signal.

**Fig-2:**
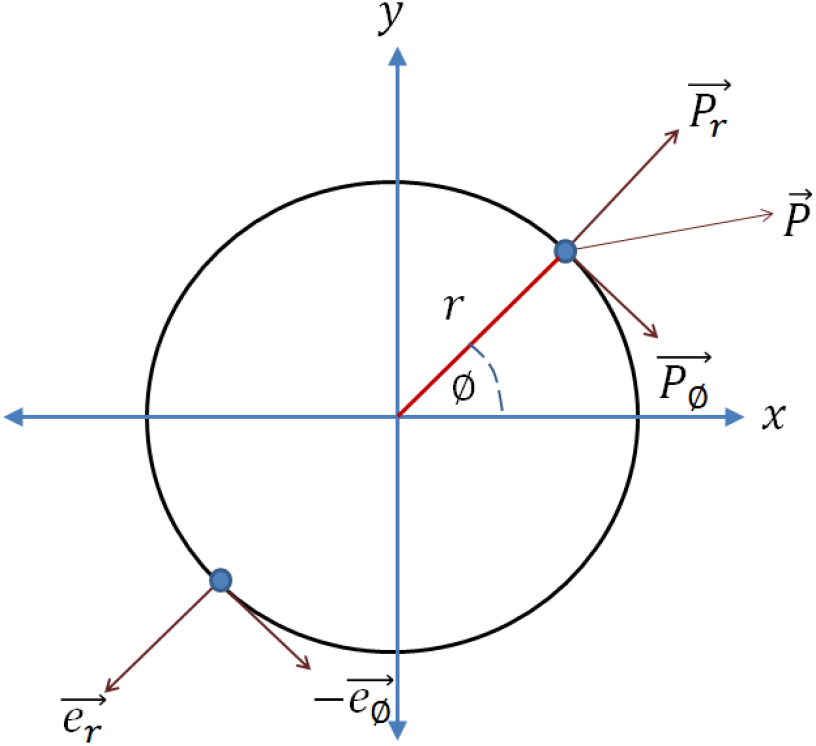
Considering the circle as the limit cycle of the Hopf oscillator, 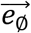 is the unit vector along increasing azimuth angle 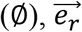 is the unit vector along radius (*r*), 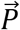 is the overall external input perturbation.

This motivates us to adapt the learning rule for the natural frequency of the oscillator, as proposed in equation-2c. We have simulated the equations-2a to 2c and observed that the proposed learning rule for the natural frequency of the oscillator allows it to adapt to the frequency of the input complex sinusoidal signal, as shown in the following figure (fig-3).

**Fig-3:**
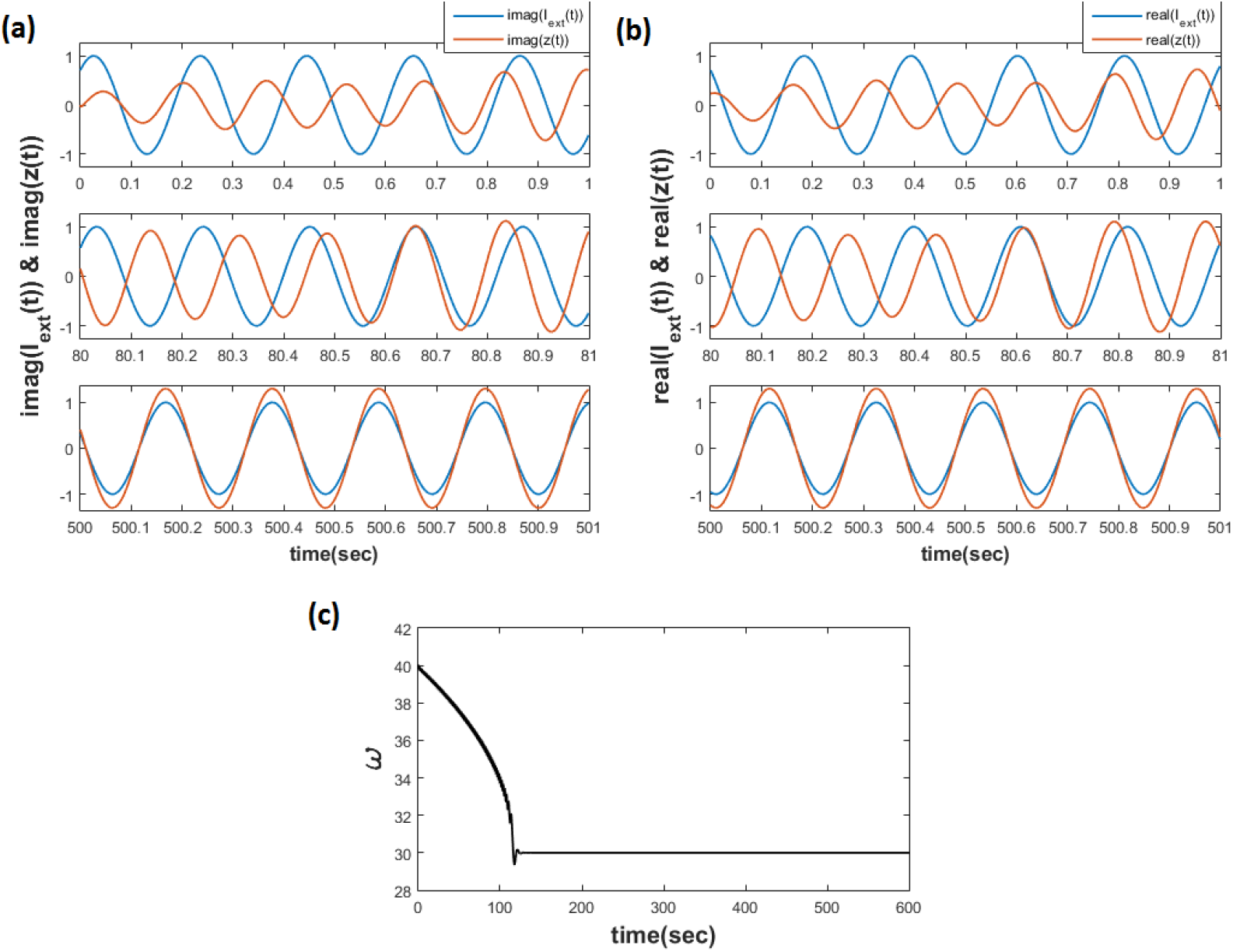
Equation-2a to 2c is simulated for *μ* = 1, *ω*(0) = 40, *ω*_0_ = 30, *ε* = 0.9, *A* = 1, 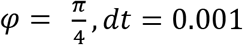 *sec* for 1000 secs. (a) Depicts the variation of *x*(*t*) w.r.t *real*(*I_ext_*(*t*)) at various time instants. (b) Depicts the variation of *y*(*t*) w.r.t *img*(*I_ext_*(*t*)) at various time instants. (c) It can be observed that the natural frequency of the Hopf oscillator adapts to the frequency of the input signal.

### 2.3 A pair of Hopf oscillators coupled through real coupling

When two Hopf oscillators with equal natural frequencies are coupled with the real coupling coefficient as described below (equation-3), they are going to phase lock in phase (0) or out of phase by (2*n* + 1)*π* depending on the polarity of the coupling and initialization. (for proof refer to appendix-A1)

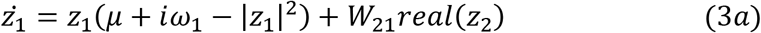

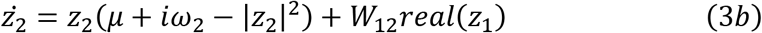

where *W*_21_ is the real coupling coefficient from 1^st^ oscillator to 2^nd^ one, and *W*_12_ is the real coupling coefficient from 2^nd^ to 1^st^. In the stated scenario, both of the oscillators have identical natural frequencies, *ω*_1_ = *ω*_2_. This, with real-valued coupling, a pair of Hopf oscillators with equal intrinsic frequencies can only produce two possible values of phase difference (fig-4).

**Fig-4:**
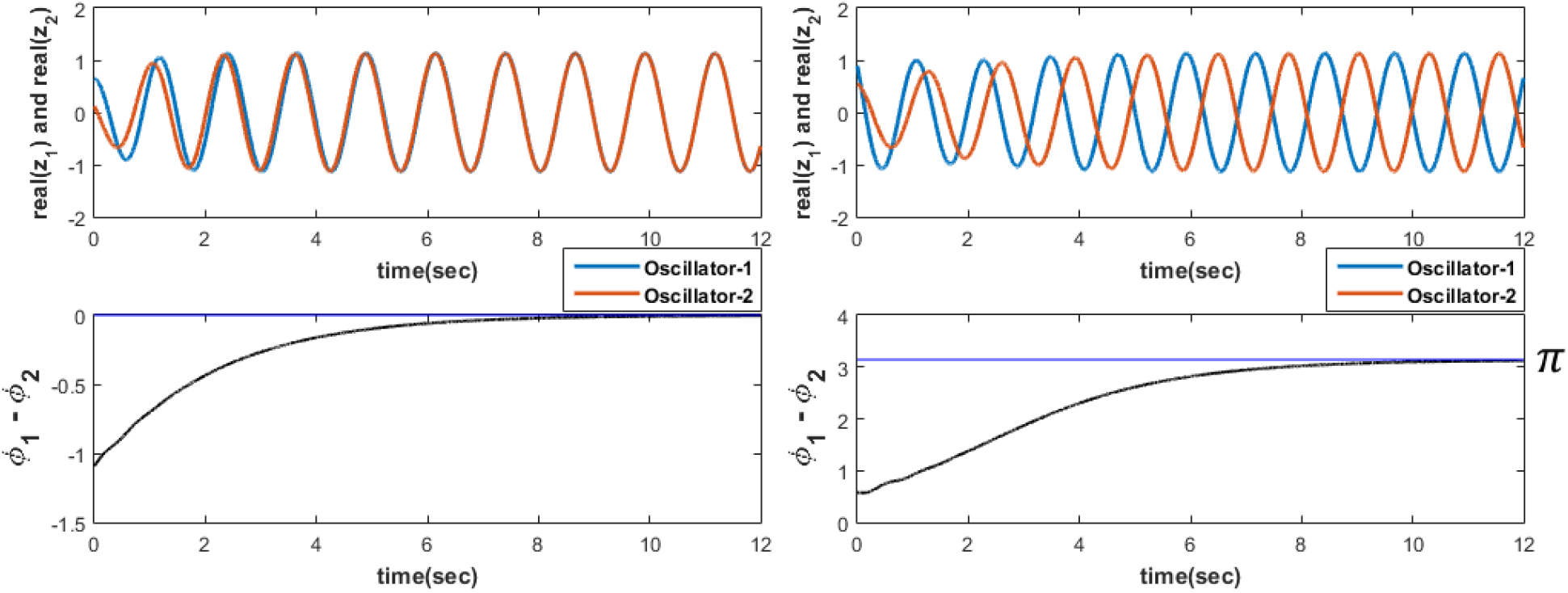
Two coupled Hopf oscillators with real coupling coefficients, as described in equation-3, are simulated under two different scenarios with *μ* = 1 and *ω*_1_ = *ω*_2_ = 5. In the 1^st^ case where *W*_21_ = *W*_12_ = 0.5, they phase lock-in phase, whereas, in the 2^nd^ one with *W*_21_ = *W*_12_ = −0.5, they phase lock 180° out of phase.

### 2.4 A pair of Hopf oscillators with Complex coupling

When two Hopf oscillators with identical natural frequencies are coupled bilaterally through complex coefficients with Hermitian symmetry, they can exhibit phase-locked oscillation at a particular angle similar to the angle of complex coupling coefficient. (for proof refer to the appendix-A2)

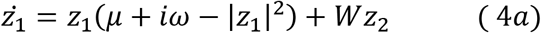

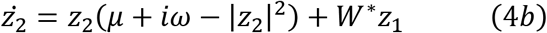

where 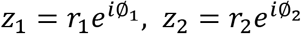 and 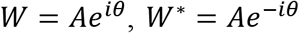 to be the coupling coefficient, in polar coordinate system representation:

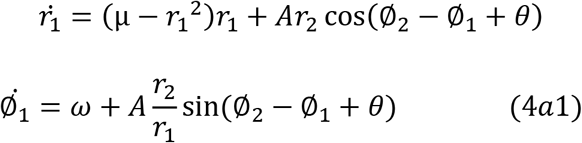

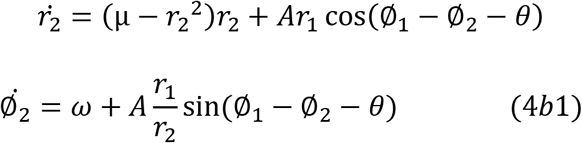

At steady-state 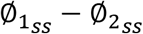 approaches any of the solutions 2*nπ* + *θ* depending on the initial conditions 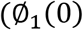 and 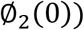, whereas the magnitude of the complex coupling coefficient determines the rate at which the phase-locking occurs. *ψ_ss_* (where *ψ* = *ϕ*_1_ – *ϕ*_2_) attains solution 2*nπ* + *θ* for the following initial condition 2*nπ* – (−*π* + *θ*) < *ϕ*_1_(0) – *ϕ*_2_(0) < 2*nπ* – (*π* + *θ*).

#### Simulated result

To verify the above result, we have simulated equation-4a1 and 4b1 numerically by Euler integration method on MATLAB platform with Δ*t* = 1 *msec*. It can be observed from the plot (refer to the figure-5) that the steady-state phase difference between the two oscillators (*ψ_ss_*) is either achieving *θ* or 2*π* + *θ* depending on the whether it was initialized in the intervals *θ* – *π* < *ψ*(0) ≤ *θ* + *π* and *θ* + *π* < *ψ*(0) ≤ 2*π* + *θ* respectively.

**Fig-5:**
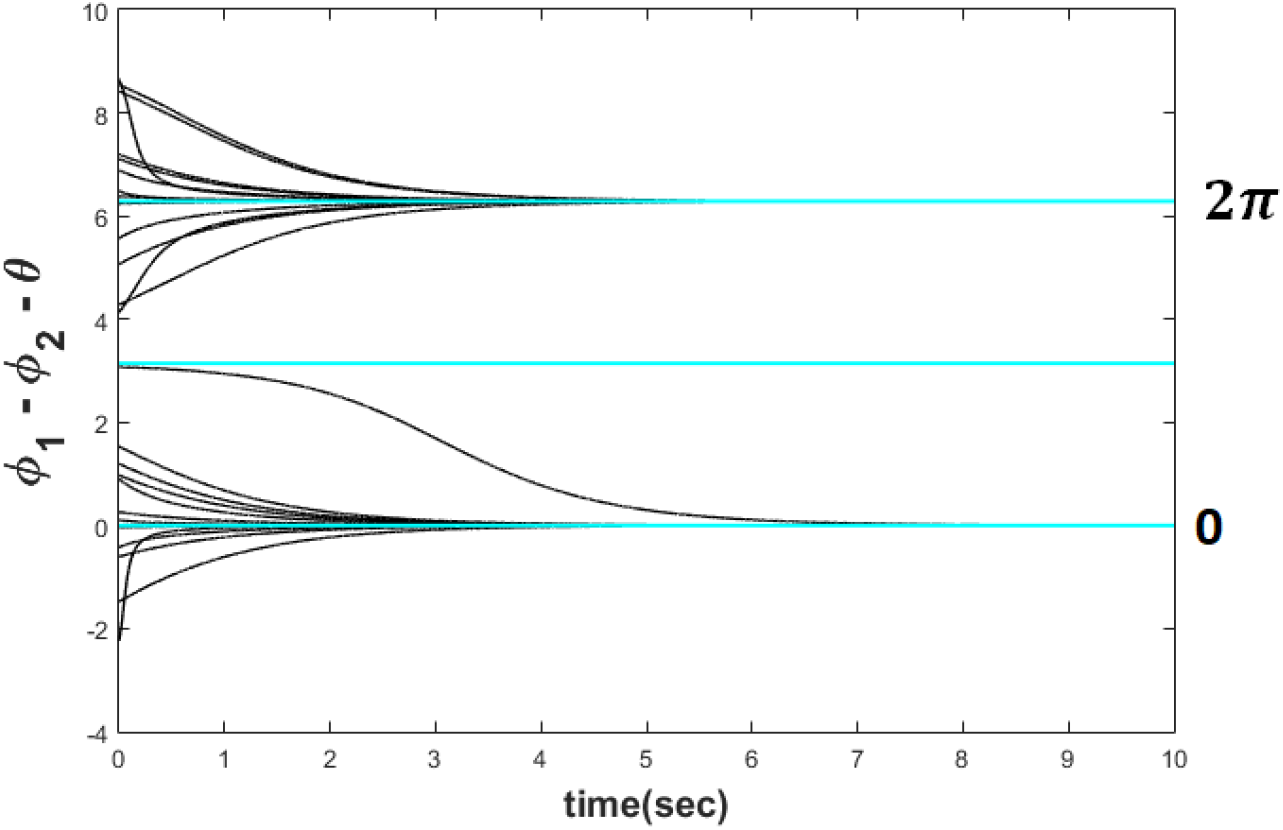
Two Hopf oscillators, dynamics of which is defined in the equation-4a1 and 4b1 simulated with 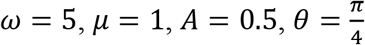 for various initial conditions (–*π* < *ϕ*_1_(0) – *ϕ*_2_(0) – *θ* ≤ 3*π*) depicting that *ψ_ss_*(= *ϕ*_1_*ss*__ – *ϕ*_2_*ss*__) can reach any of the following solutions 2*nπ* + *θ* depending on the initial condition.

### 2.5 Training rule for the complex coupling

When a pair of complex sinusoidal inputs with identical frequency is presented to a pair of coupled Hopf oscillators with complex coupling and identical natural frequencies (equation-4), and when the complex coupling coefficient is adapted according to equation-5, the angle of the coupling coefficient approaches the phase difference of the external inputs. In other words, the complex coupling learns the phase relationship of the external inputs.

To train the complex coupling coefficient, a Hebbian like learning rule can be used as follows:

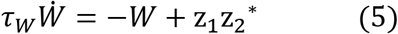

Assuming *τ_w_* is the time constant for the specified learning dynamics. A similar adaptation of the Hebb’s rule was used earlier to train connections in a complex version of the Hopfield network (Hopfield, 1982). When the dynamics of two Hopf oscillators is defined by equation-4 (with no external input), and the complex coupling coefficient is trained according to equation-5 with very small fixed value of *A* (|*W*|), *θ*(*angle*(*W*)) learns 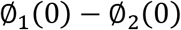. From equation-4, it can be interpreted that 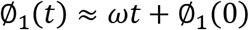 and 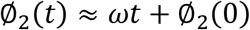 (*Assuming A* ≪ 1), similarly from equation-5, we can see that *θ or angle*(*W*) learns *angle*(*z*_1_*z*_2_*) or 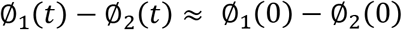.

On the other hand, in a network of two coupled oscillators driven by separate external sinusoidal forcing as described in the following equation (eq-6) with the same frequency as the natural frequency of the two oscillators but any initial phase offset (*φ*_1_,*φ*_2_) will drive those oscillators to oscillate at the same phase as the external sinusoidal forcing (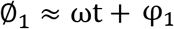 *and* 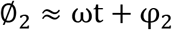), provided very low magnitude of the complex coupling coefficient (*A* ≪ 1).

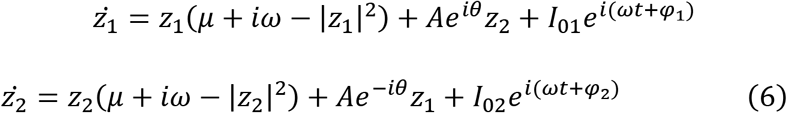

Under the influence of external input, the angles of *z*_1_ and *z*_2_ (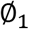 and 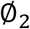 respectively) tend to ωt + φ_1_ and ωt + φ_2_, respectively. When the complex coupling coefficient is trained according to equation-5, the *angle* (*W*) or *θ* learns *angle*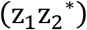 or 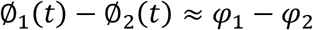. During training, A must be kept low (*A* ≪ 1 of the order of 10^-4^) so that z_1_ and z_2_ dynamics is influenced only by the intrinsic oscillatory dynamics and the forcing signal. During testing, however, the external input is removed, A is again increased (at the order of 10^-2^ and above) so that the phase difference between the oscillators is stable and determined by the coupling constant.

#### Simulated result

The proposed Hebb like learning rule (equation-5) allows the angle of complex coupling weight (*θ*) to learn the phase difference between the complex sinusoidal inputs driving each of the oscillators for very low fixed magnitude of the complex coupling weight (*A* ≪ 1). To verify this, equation-6 is simulated for the set of parameters as described in the following figure (fig-6). It can be observed that *θ* learns the phase difference between the complex sinusoidal input signals (*φ*_1_ – *φ*_2_).

**Fig-6:**
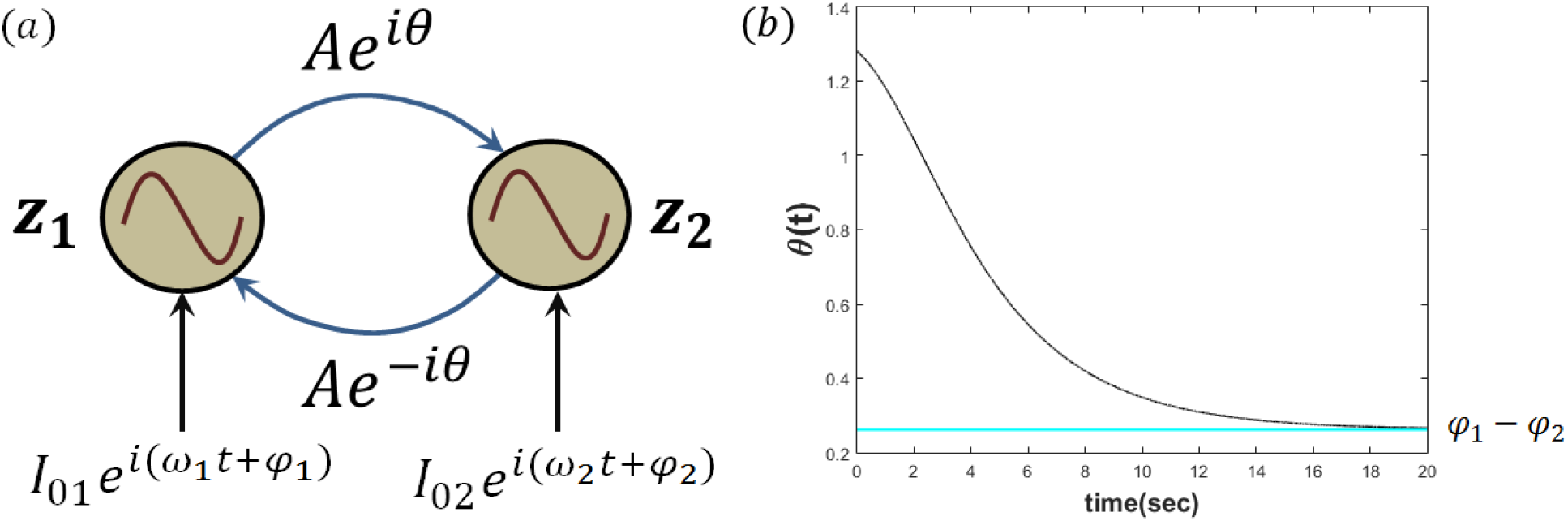
(a) Network schematic of dynamics represented in equation-6 where the complex coupling coefficient is trained according to equation-5. (b) It can be verified that for the following set of network parameters *θ* learns 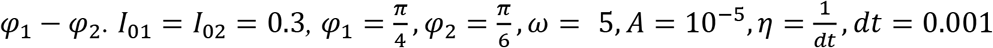.

### 2.6 Coupling two Hopf oscillators with different natural frequencies

A different kind of dynamics can be observed when two sinusoidal oscillators with different natural frequencies are coupled with real coupling coefficient. A pair of Hopf oscillators with real coupling shows Arnold tongue behavior which may be described as follows: the two oscillators entrain to commensurate frequencies (having a simple integral ratio) for specific ranges of the coupling strength, and the range of frequencies for which the entrainment occurs widens with the strength of the coupling coefficient (Izhikevich). There are whole ranges of the coupling strength and the natural frequencies where there is no specific phase relationship at all. Therefore, it is not easy to get a stable phase relationship between two Hopf oscillators with very different frequencies and real coupling.

The situation is a bit more facile with Kuramoto oscillators (Izhikevich). When a pair of Kuramoto oscillators are coupled according to the equation shown below, their natural frequencies converge to a particular frequency in between their natural frequencies, depending on the coupling strength.

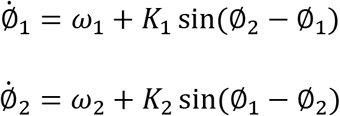

Let, *K*_1_ and *K*_2_ be the real coupling strengths in the forward and backward directions, respectively. The phase difference between the two oscillators 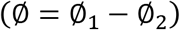 reaches a steady-state value if *K*_1_ + *K*_2_ > |*ω*_1_ – *ω*_2_| which is satisfied by the relationship 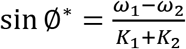 where 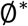 be the steady-state phase difference. Notably, both of the oscillators reach a periodic solution with frequency 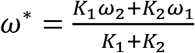. In this scenario, although synchronization at a particular intermediate frequency with a particular phase difference is ensured, the natural frequency of oscillation is not maintained, i.e., to phase lock at a particular angle the oscillation of both the oscillators had to converge at a particular intermediate value. A network of such Kuramoto oscillators (equation-7a) with natural frequencies drawn from a Gaussian distribution with constrained standard deviation can show increased phase synchrony among the oscillators when the isotopic coupling strength (K) exceeds a threshold (Breakspear et al., 2010).

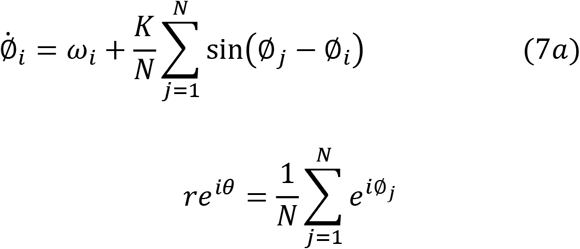

Thus, we can see that the dynamics of a pair of coupled Hopf oscillators become dramatically more complicated when we relax the equality relationship between the natural frequencies of the coupled oscillators. We can visualize a simple, desirable extension of the dynamics of a pair of coupled oscillators with distinct natural frequencies. When the natural frequencies are equal, we found that the phase difference equals the angle of the complex coupling factor. However, when the natural frequencies are unequal, there cannot be a phase difference in the usual sense. Therefore, one may define a quantity called “normalized phase difference” as follows:

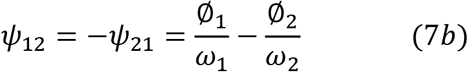

Let *ψ*_12_ and *ψ*_21_ be the normalized phase difference of 1^st^ oscillator w.r.t 2^nd^ oscillator and vice versa. Is it possible to relate the normalized phase difference with the angle of the complex coupling coefficient? It turns out that such a relationship is not possible even with the complex coupling of equation-4. To this end, we extend the complex coupling to a new form of coupling we label “power coupling” as described below.

### 2.7 “Power coupling” between a pair of Hopf oscillator

A pair of sinusoidal oscillators (Hopf or Kuramoto oscillator) can entrain at a specific normalized phase difference if they are coupled through complex “power coupling” coefficient according to the following equation (equation-8a and 8b) at a particular value dependent on the angle of complex “power coupling” coefficient, natural frequencies of the coupled oscillators and the initial values of the phase angle of those oscillators (for proof please refer to appendix-A3).

Coupling a pair of Hopf oscillators through *power coupling*:

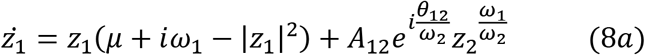

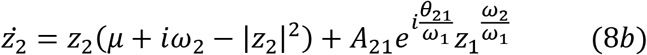

where, 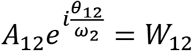 is the weight of *power coupling* from 2^nd^ oscillator to the 1^st^ oscillator and 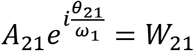 is the weight of *power coupling* from 1^st^ oscillator to the 2^nd^ oscillator.

The polar coordinate representation:

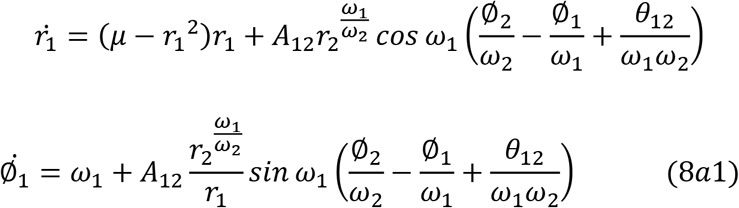

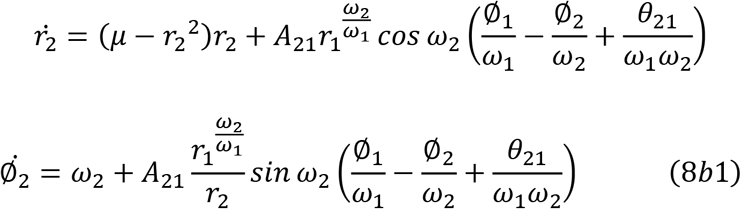

The schematic of the coupling architecture is elaborated in the following figure (fig-7).

**Fig-7:**
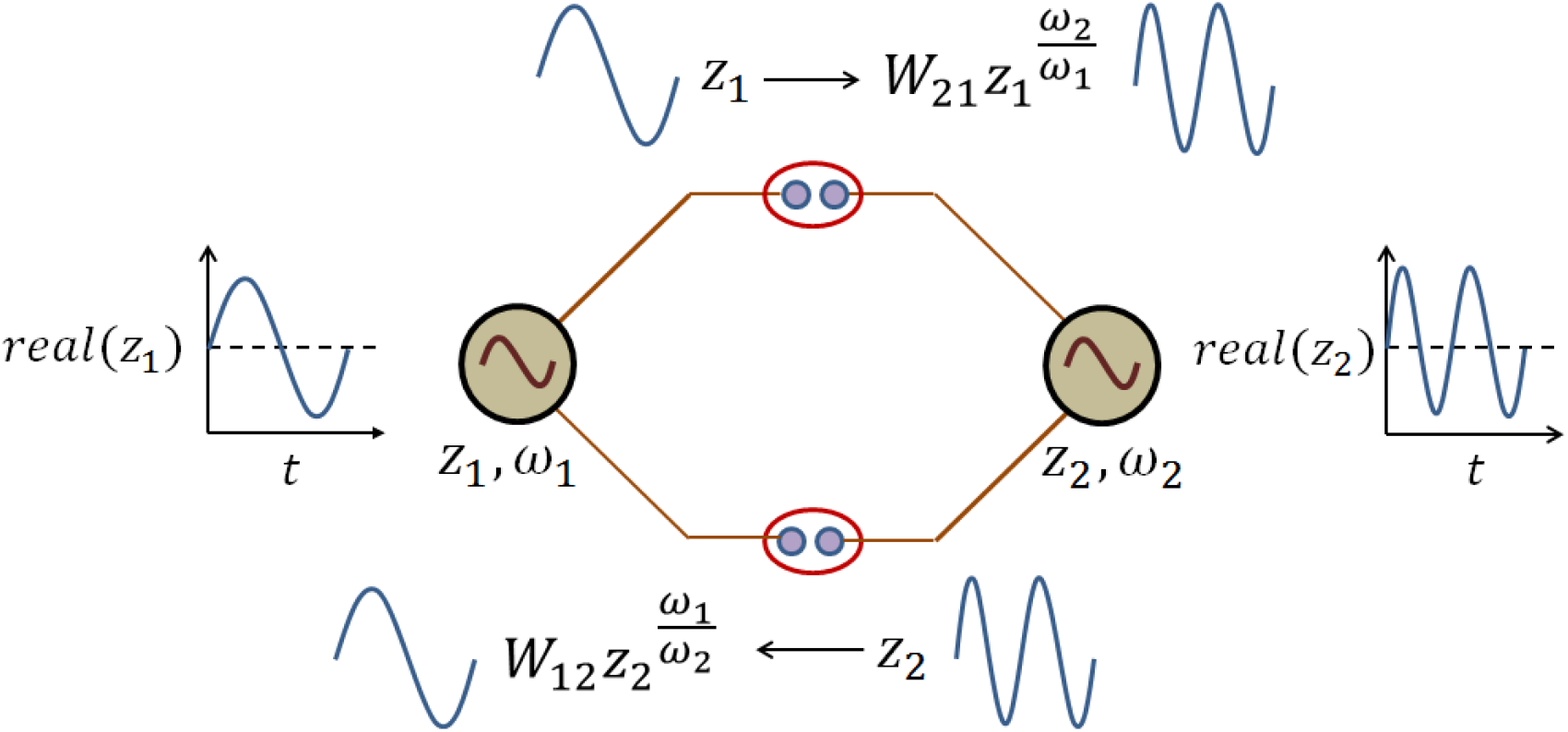
The schematic network representation of equation-8. There is a kind of frequency transformation occurring at the *power coupling* synapse because of the following term in the dynamical equation-8: 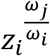. At the *power coupling* synapse, this frequency-transformed version of the oscillation from *j^th^* oscillator is weighted by the power coupling weight 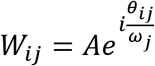 to convert the forcing signal from *j^th^* oscillator to *i^th^* oscillator and vice versa.

Based on equations-8a1 and 8a2, we understand that two modified Kuramoto oscillators may be coupled using power coupling as follows:

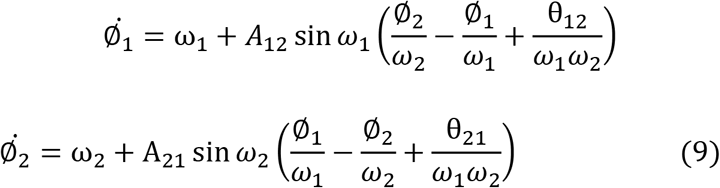

### 2.8 Power coupling among N Hopf oscillators

Similarly, the dynamics of *the i^th^* oscillator in a network of *N* supercritical Hopf oscillators coupled through *power coupling* can be represented as:

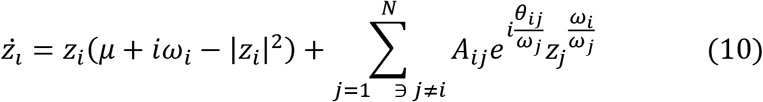

Let 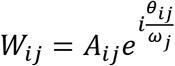 be the weight of power coupling from the *j^th^* oscillator to *i^th^* oscillator.

The polar coordinate representation is:

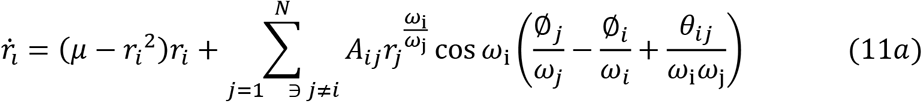

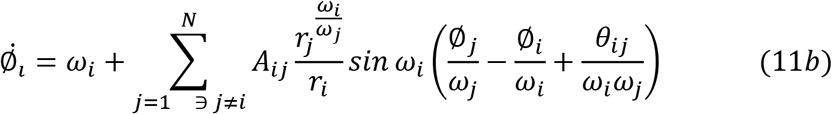

There fore, the dynamics of *the i^th^* oscillator in a network of *N* number Kuramoto oscillators coupled through power coupling can be represented as:

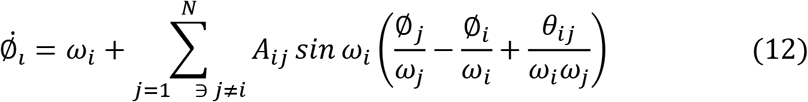

We may appreciate the difference in the second term in equation-7 and equation-12. 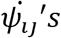 (defined in equation-7b) can be obtained from equation-8a1, 8b1 and 11b for a network of 2 oscillators and also for the general case of *N*(> 2) Hopf oscillators respectively:

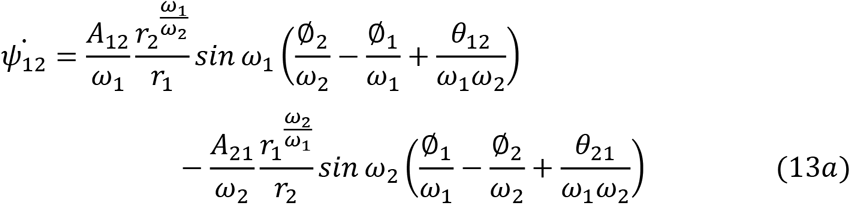

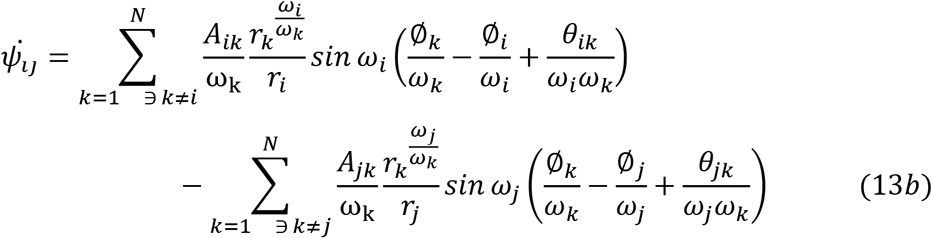

At steady state, *ψ*_12_ and 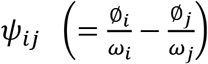 can attain any of the possible solutions of the above equations (equation-13a and 13b).

For a network of any arbitrary *N* number of Hopf oscillators as described in equation-10 and 11 under the constraint 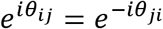 or 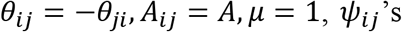 can achieve the following *desirable* solution:

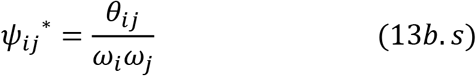

for certain initial conditions.

However, there can exist other solutions for which the sum of the terms on the right-hand side of equation-13a and 13b is zero, without all the individual terms being zero. That is equation-13b.s above is not satisfied. We call these solutions *spurious* solutions.

Whether the final solution is a desirable or a spurious solution depends on the initial conditions. Therefore, the solution *ψ_ij_*\* can be achieved only for certain initial values of 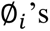.

#### Two Hopf oscillators with power coupling

Under the following constraints *θ*_12_ = *θ, θ*_21_ = – *θ,A*_12_ = *A*_21_ = *A, μ* = 1, some of the steadystate solutions of equation-13a where both entities on the right-hand side of equation-13a is zero, is as follows:

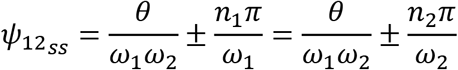

following the specification: 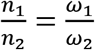. It can be noticed that *n*_1_ = *n*_2_ = 0 gives us the desired solution, as mentioned in equation-13b.s.

Let 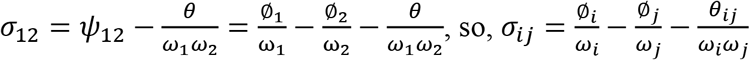.

From the following figure (fig-8) it can be verified that *ψ*_12_ and *σ*_12_ attain 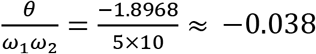 and zero solution (the desired solution as mentioned before) respectively for the initialization specified in the figure caption.

**Fig-8:**
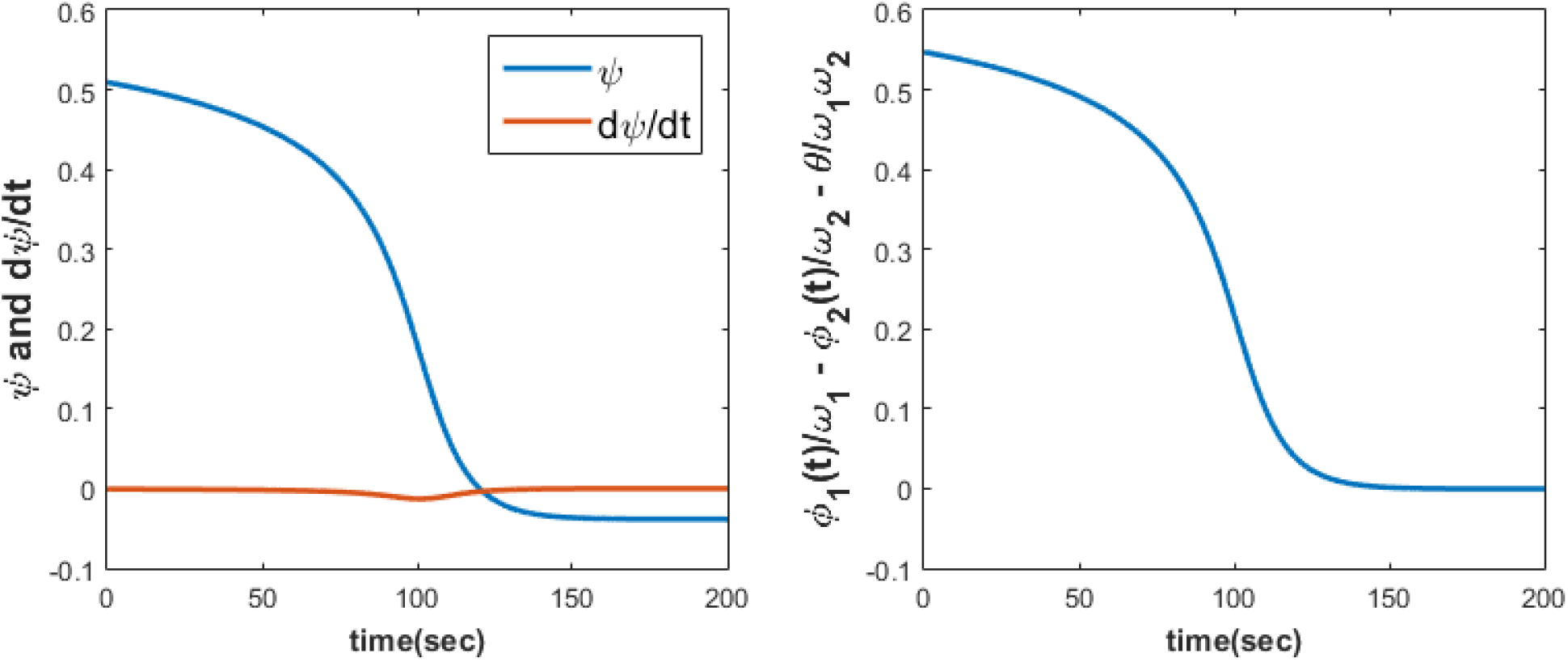
The variation of 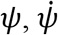, and 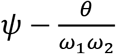 w.r.t time is plotted by simulating equation-8a1 and 8b1 for the following set of parameters and initial conditions. 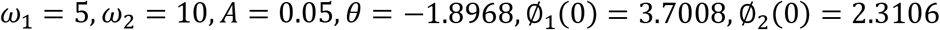.

To check the dependency of *σ*_12_*ss*__ on the initial values of 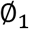 and 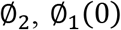 and 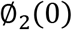 space is discretized at 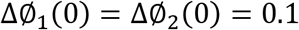, in the interval 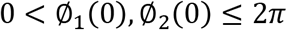 and equation-8a1 and 8b1 is simulated for 200 secs to ensure the steady-state to be achieved for various combinations of 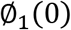 and 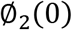. From the following figure (fig-9) it can be observed that *σ*_12_*ss*__ achieves the following solutions: 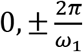; for *n*_1_ = 0,1 or 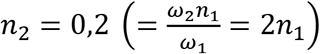 independent of *θ*.

**Fig-9:**
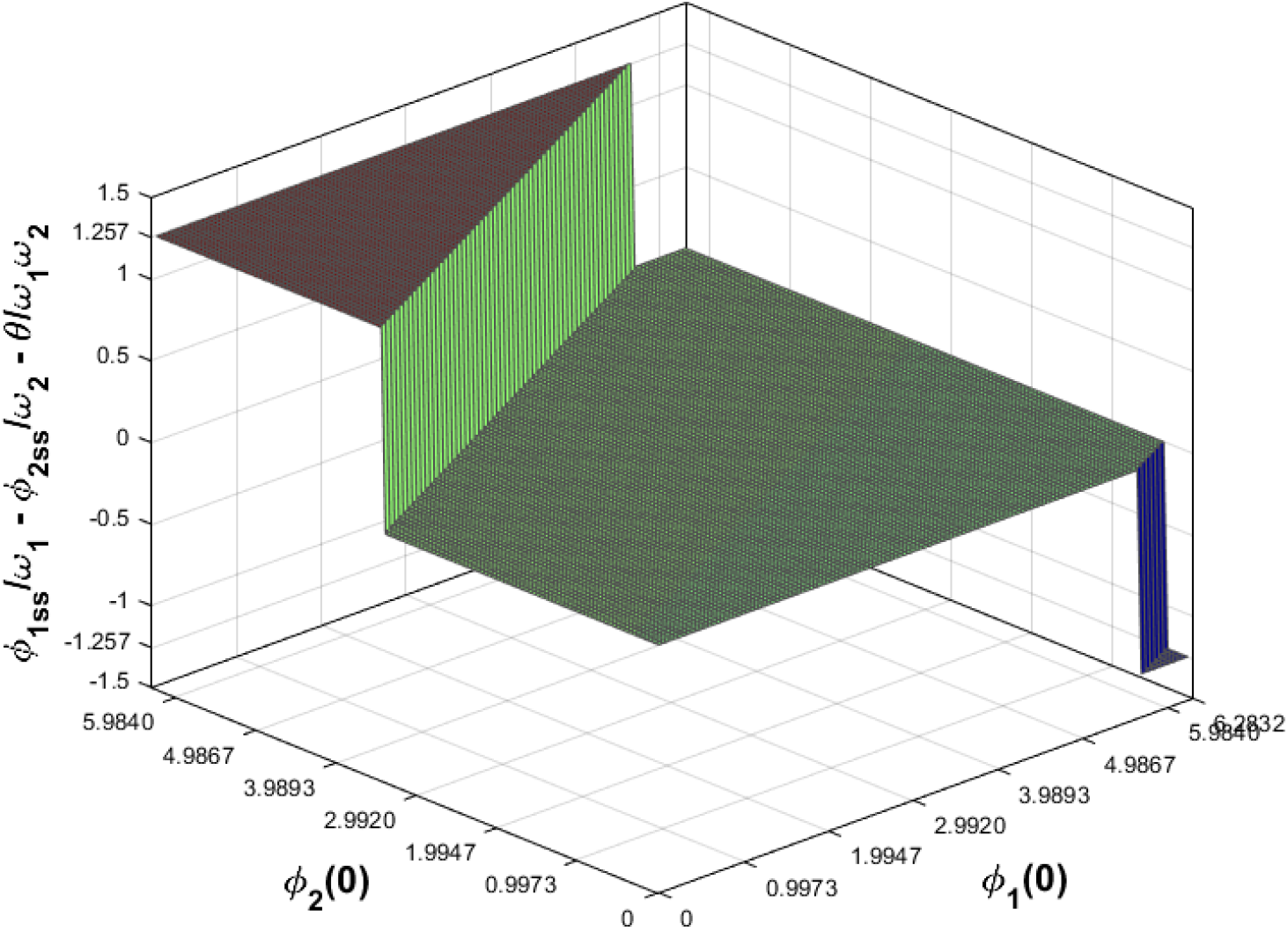
It can be verified that depending on the various combinations of initial conditions 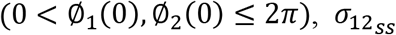 can reach one of the solutions 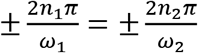. For the following set of parameters, simulated results show that *σ*_12_*ss*__ reaches one of the following three solutions 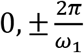 *or* 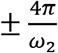 *or* ± 1.257. Simulation parameters: *ω*_1_ = 5, *ω*_2_ = 10, *A* = 0.2, *θ* = 2.9644.

#### Three Hopf oscillators coupled through power coupling

We now explore the solution space numerically for a three-oscillator system. When three Hopf oscillators are coupled according to the equation-10 or 11 (*N* = 3) with *θ_ij_*’s, chosen as *θ_ij_* = *θ_i_* – ∀0 < *θ_i_, θ_j_* ≤ 2*π* (i.e., *θ_ij_* = –*θ_ij_*), at steady state *σ_ij_* achieves one of the possible solutions depending on the initializations 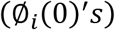. The 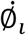 dynamics of the three oscillators can be expressed in the following form.

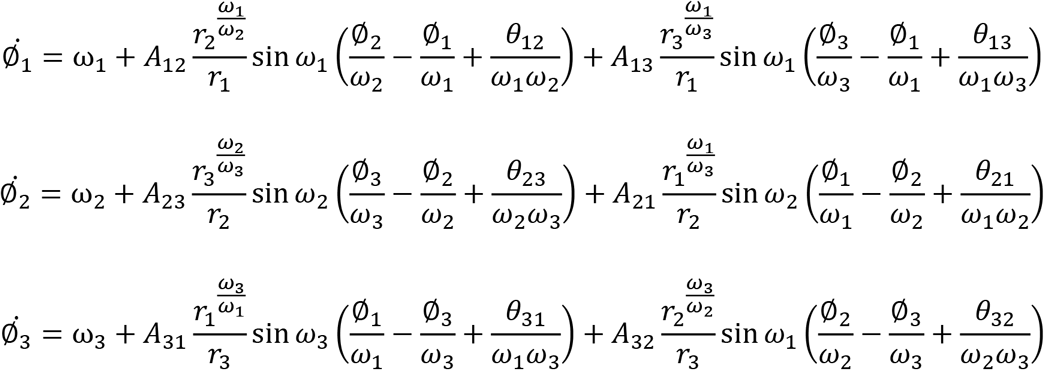

At steady-state 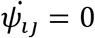, where 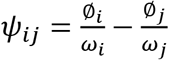 or 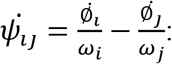 Assuming *μ* = 1, *A_ij_* = *A* and *e^iθ_ij_^* is a Hermitian matrix (*i.e*., 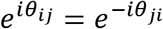),

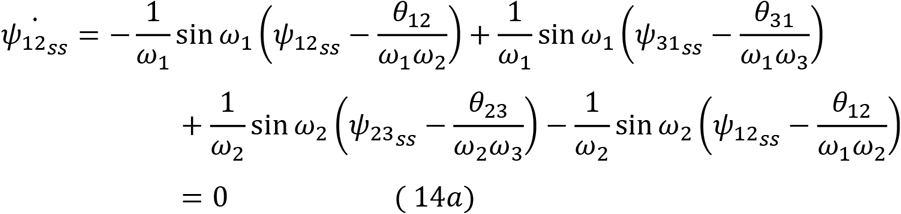

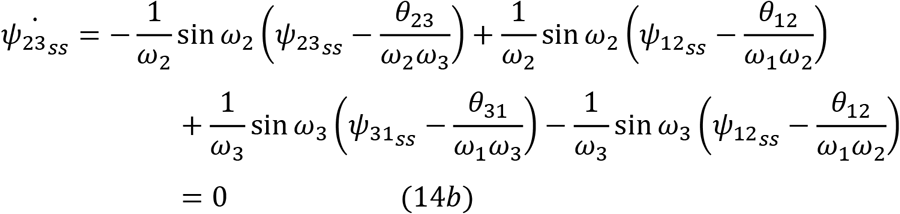

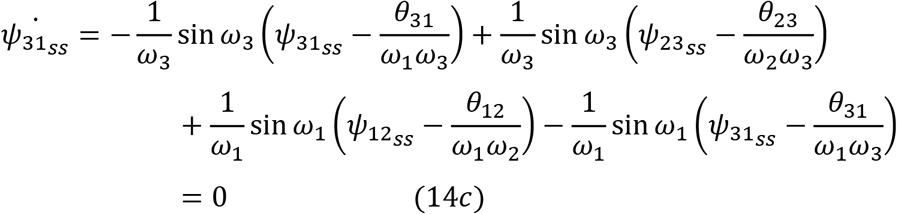

The trivial and desirable solution set for the above equation is:

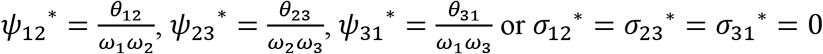

However, like before in equation-(13a), there are several spurious solutions for which the entire right-hand sides of equations-(14a, 14b, 14c) are zero, without each of the individual terms being zero separately. So, *σ_ij_* can achieve any of the possible solutions satisfying the above equation. It appears that finding the spurious solutions analytically are not possible.

To verify this numerically equation-11 was simulated for the given set of network parameters: 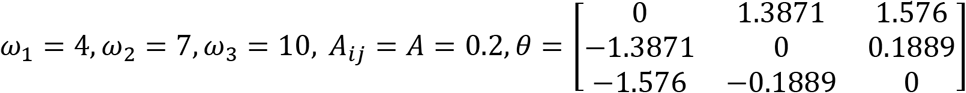 and the initial values of 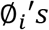 in the range 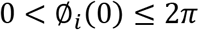 with 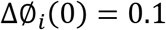 for 100 secs to let the system achieve a steady-state. It can be observed from equation-14 that the possible solutions of *σ_ij_ss__* do not depend on *θ_ij_* but on the natural frequencies of the oscillators (*ω_i_*). The following figure (fig-10) summarizes the dependency of *σ_ij_ss__* on 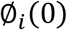 and 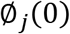 where it can be observed that *σ*_12_*ss*__, *σ*_23_*ss*__, and *σ*_31_*ss*__ achieve any of the 5, 3, 4 solutions respectively (mentioned in the figure caption) and particular combinations of these solutions satisfy the above equation. For a given initial value of 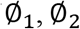 and 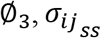’s attained one of the solutions and for a confined subspace of 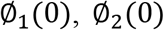 and 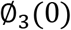 under space 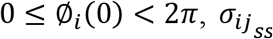’s attained the zero or desired solution (*σ_ij_*\*).

**Fig-10:**
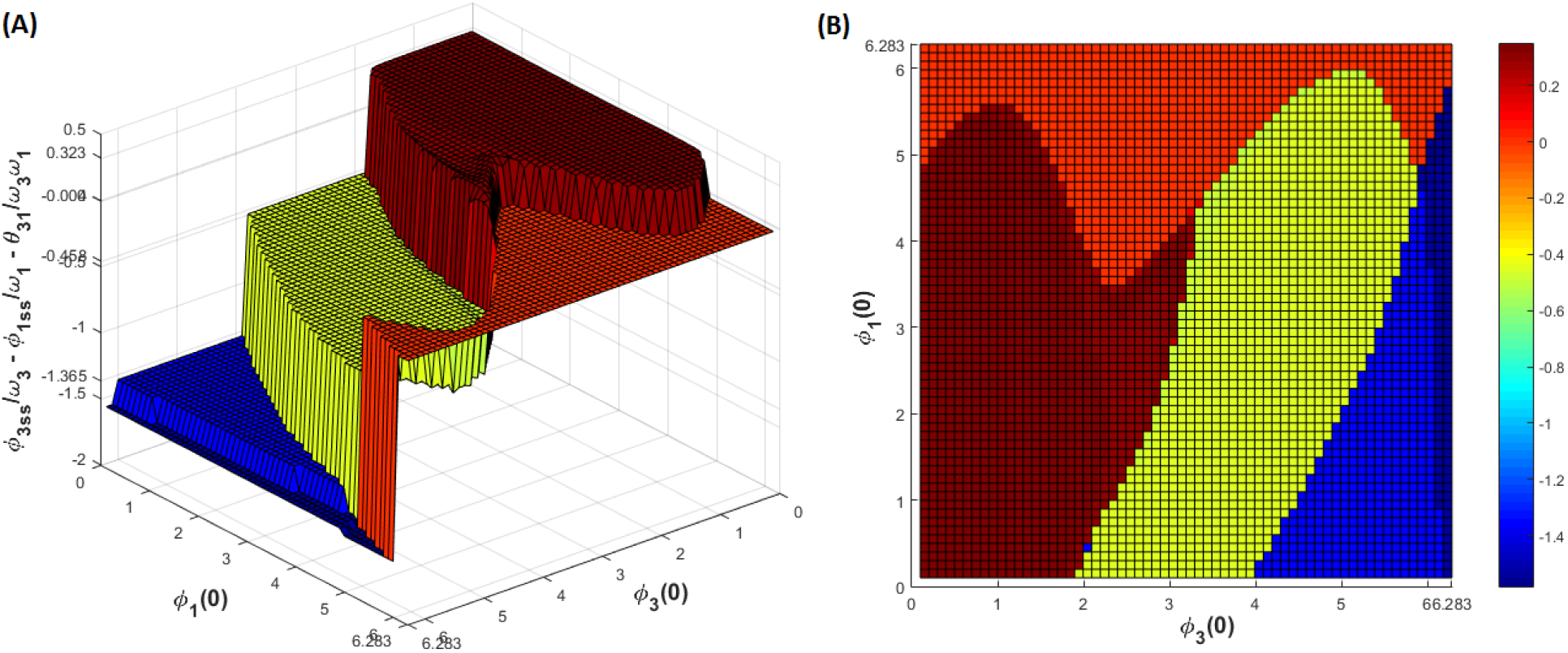
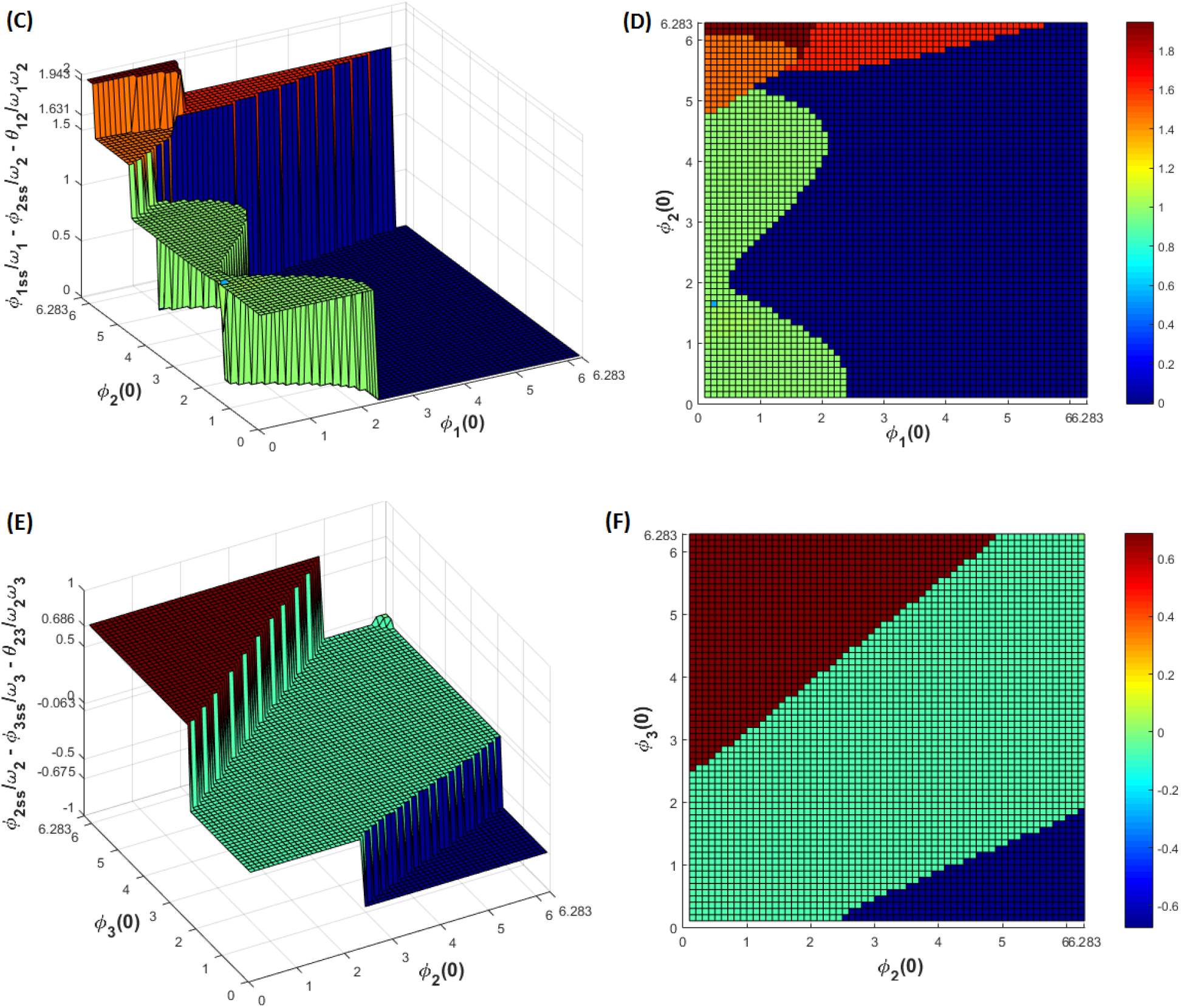
Equation-11 is simulated for 100 secs for the network parameter values mentioned previously to check for what combinations of initial values of 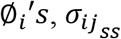’s reaches, which of the solutions of equation-14. *(a)nd (b)* depicts that for various initial values of 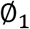 and 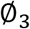 in between 0 *to* 2*π*, *σ*_31_*ss*__ attains 4 possible solutions: 0.3235, –0.0036, –0.4576, –1.3652, similarly from (*c*), (*d*) and (*e*), (*f*) it can be seen that for various initial values of 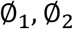 and 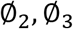 in between 0 *to 2π, σ*_12_*ss*__ and *σ*_23_*ss*__ attains the 0.99,1.465,1.6307,1.9426, —0.0046 and 0.0633, 0.6855, –0.6751 solutions respectively, satisfying equation-14.

### 2.9 Hebbian learning for the N-oscillator system with power coupling

We may extend the Hebbian learning of complex coupling (eqn. 5), to the case of power coupling as follows.

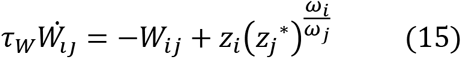

Let *τ_w_* be the time constant of the above learning dynamics. It is shown both analytically and numerically that *ω_j_* times *angle* (*W_ij_*) or *θ_ij_* tries to learn 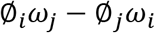 when the weight of the *power coupling* is trained according to equation-15 (for proof, please refer to appendix-A4). Similarly, as it was previously shown during the training or phase encoding of the complex coupling coefficient with very small fixed value of *A_ij_* (magnitude of complex coupling coefficient), *θ_ij_* (angle of complex coupling coefficient) learns 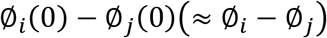 for arbitrary values of *ω* (constraining the natural frequencies to be identical). When the *power coupling* weight is updated according to equation-15 under the same constraint as *A_ij_* ≪ 1 and 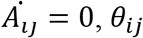 learns 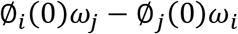 when there is no external input.

From equation-11, it can be interpreted that 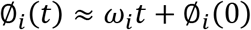 for *A_ij_* ≪ 1. So, when a network of sinusoidal oscillators coupled according to equation-10 or 14 with the power coupling weights 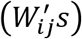 trained according to equation-15, *θ_ij_* learns 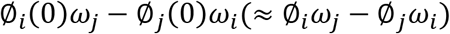.

In such a network when individual oscillators (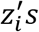 or 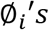) are driven by complex sinusoidal forcing (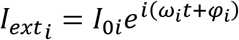 *for the i^th^ oscillator*) and the power coupling weights are trained as stated, *θ_ij_* learns *ψ_i_ ω_j_* – *φ_j_ω_i_* as with 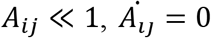 and 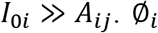’s are approximately same as *ω_i_t* + *φ_i_*. The dynamics of such a network of supercritical Hopf oscillators is defined below:

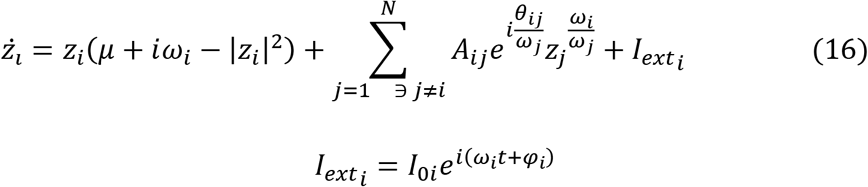

Polar coordinate representation:

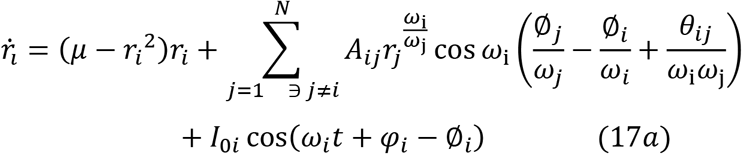

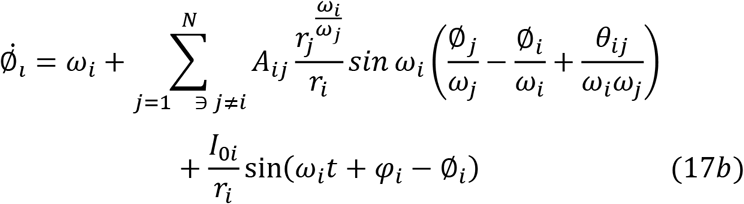

The dynamics of such a network of Kuramoto oscillators:

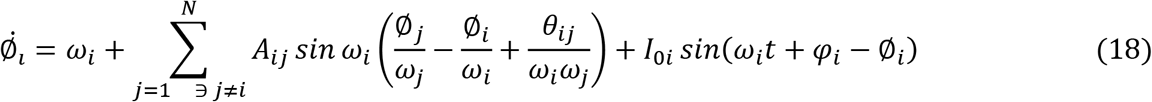

#### Simulated result

When two Hopf oscillators are coupled through power coupling with very weak coupling coefficient (*A_ij_* ≪ 1) as described above in equation-16, *θ_ij_* learns 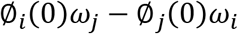 while there is no external input (*I_ext_i__* = 0) and *θ_ij_* learns *φ_i_ω_j_* – *φ_j_ω_i_* when *i^th^* oscillator is forced by complex sinusoidal external perturbation. The following simulation results support the stated argument (figs-11 and 12).

**Fig-11:**
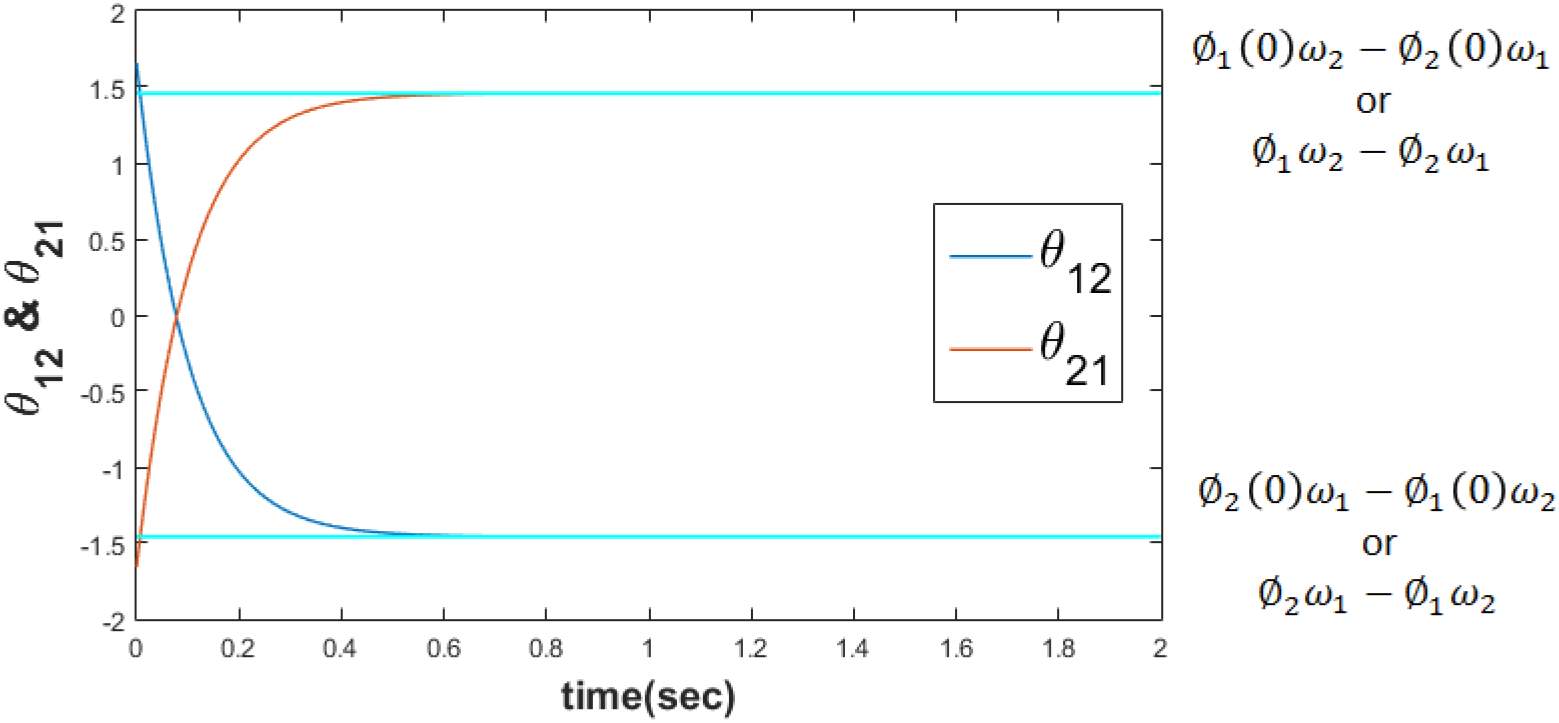
When two Hopf oscillators (*z*_1_, *z*_2_) coupled through power coupling according to equation-16 (for N=2 and *I_ext_i__* = 0) with the parameters *ω*_1_ = 5, *ω*_2_ = 10, *A*_12_ = *A*_21_ = 0.0001, *θ*_12_(0) = 1.657, *θ*_21_(0) = –1.657 and the power coupling weights are trained according to equation-15, keeping the magnitude of the power coupling weight (*A_ij_*) constant, *ω_j_* times the angle of power coupling, *θ_ij_* learns 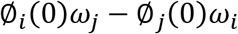 or 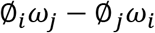. Here 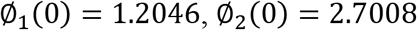, and *τ_W_* = 10^3^. It is clear from the plot that *θ*_12_ reaches 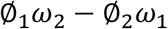 and *θ*_21_ reaches 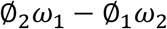 w.r.t time.

**Fig-12:**
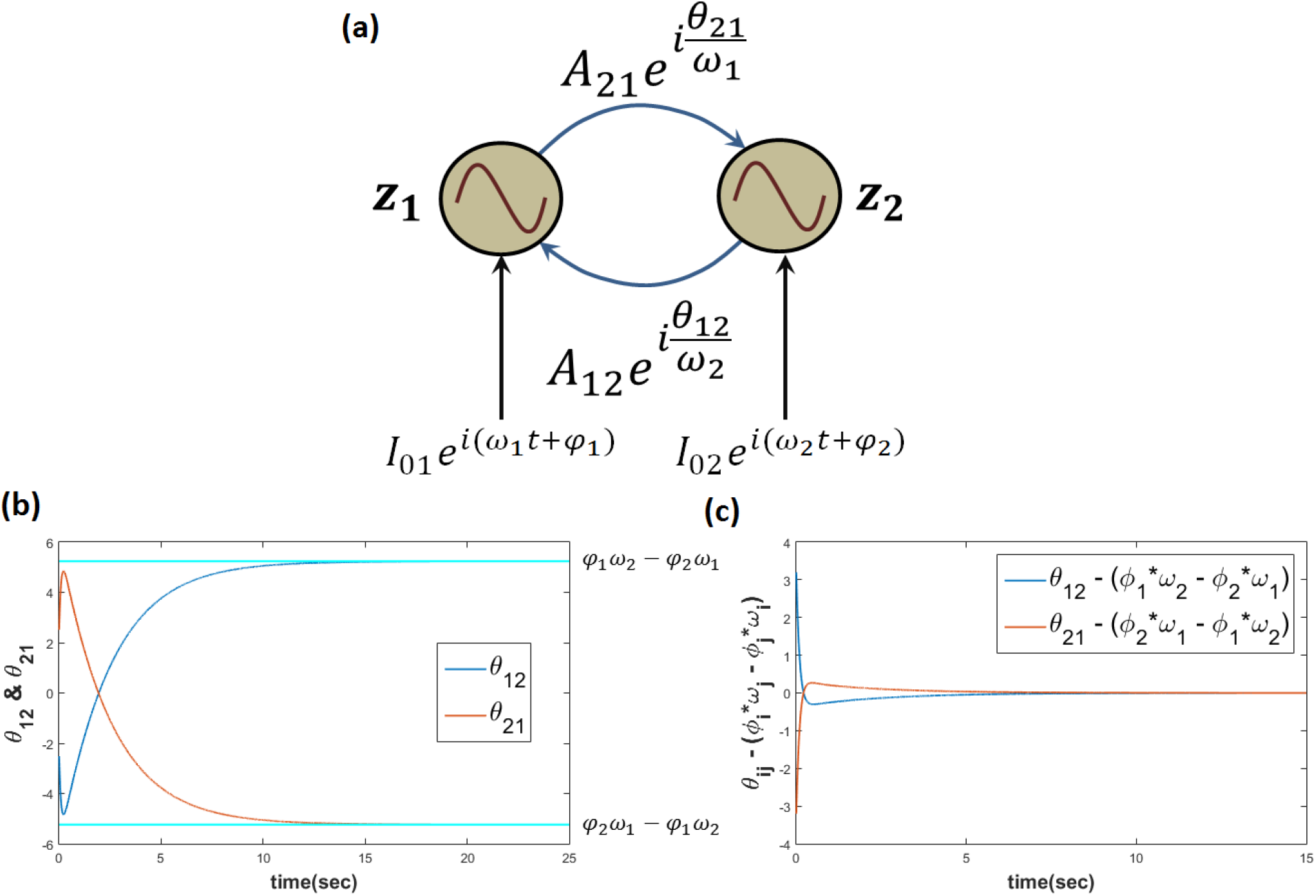
When two such Hopf oscillators as described in the previous simulation are forced by an external complex sinusoidal perturbation as described in equation-16 (*I_ext_i__* ≠ 0), the schematic of the network is elaborated in-(a), with the following parameters *ω*_1_ = 5, *ω*_2_ = 10, *A*_12_ = *A*_21_ = 0.0001, *θ*_12_(0) = —2.513, *θ*_21_(0) = 2.513,*τ_W_* = 10^3^ and 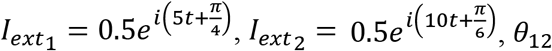, and *θ*_21_ learn *φ*_1_*ω*_2_ – *φ*_2_*ω*_1_ = –5.7131 and *φ*_2_*ω*_1_ – *φ*_1_*ω*_2_ = 5.7131 respectively (b). (c) It can be verified that the difference between *θ*_12_ and 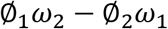 as well as *θ*_21_ and 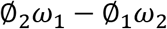 becomes zero w.r.t time.

### 2.10 A network of oscillators with adaptive frequency and trainable lateral connections

A pair of coupled adaptive Hopf oscillators driven by distinct complex sinusoidal inputs are capable of adapting their natural frequencies to the frequencies of the complex sinusoidal input signals. The trainable power coupling weight can encode the normalized phase difference between the two complex sinusoidal input signals.

When a pair of Hopf oscillators coupled through trainable power coupling coefficient (equation-15) are driven by complex sinusoidal inputs (equation-16), they adapt their natural frequencies according to equation-2b or 2c, *ω_j_* times the angle of power coupling coefficient *W_ij_* (power coupling weight from *j^th^* oscillator to *i^th^* oscillator) or *θ_ij_* can learn the normalized phase difference between the complex sinusoidal input signal. If *I_ext_i__* is the complex sinusoidal input signal driving *i^th^* oscillator, then the normalized phase difference among them is:

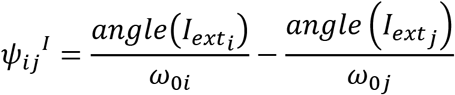

When the following dynamics is simulated for N =2 it can be verified that *θ_ij_* learns *φ_j_ω*_0*i*_ – *φ_i_ω*_0*j*_ after *ω_i_*, the natural frequency of *i^th^* oscillator learns the frequency of the input signal *ω*_0*i*_ (described in figure-13).

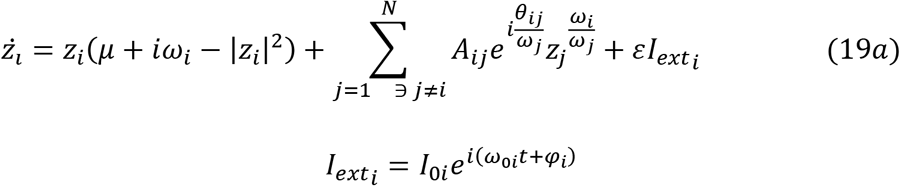

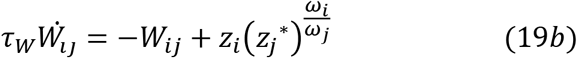

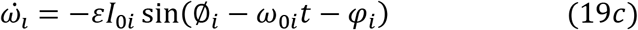

**Fig-13:**
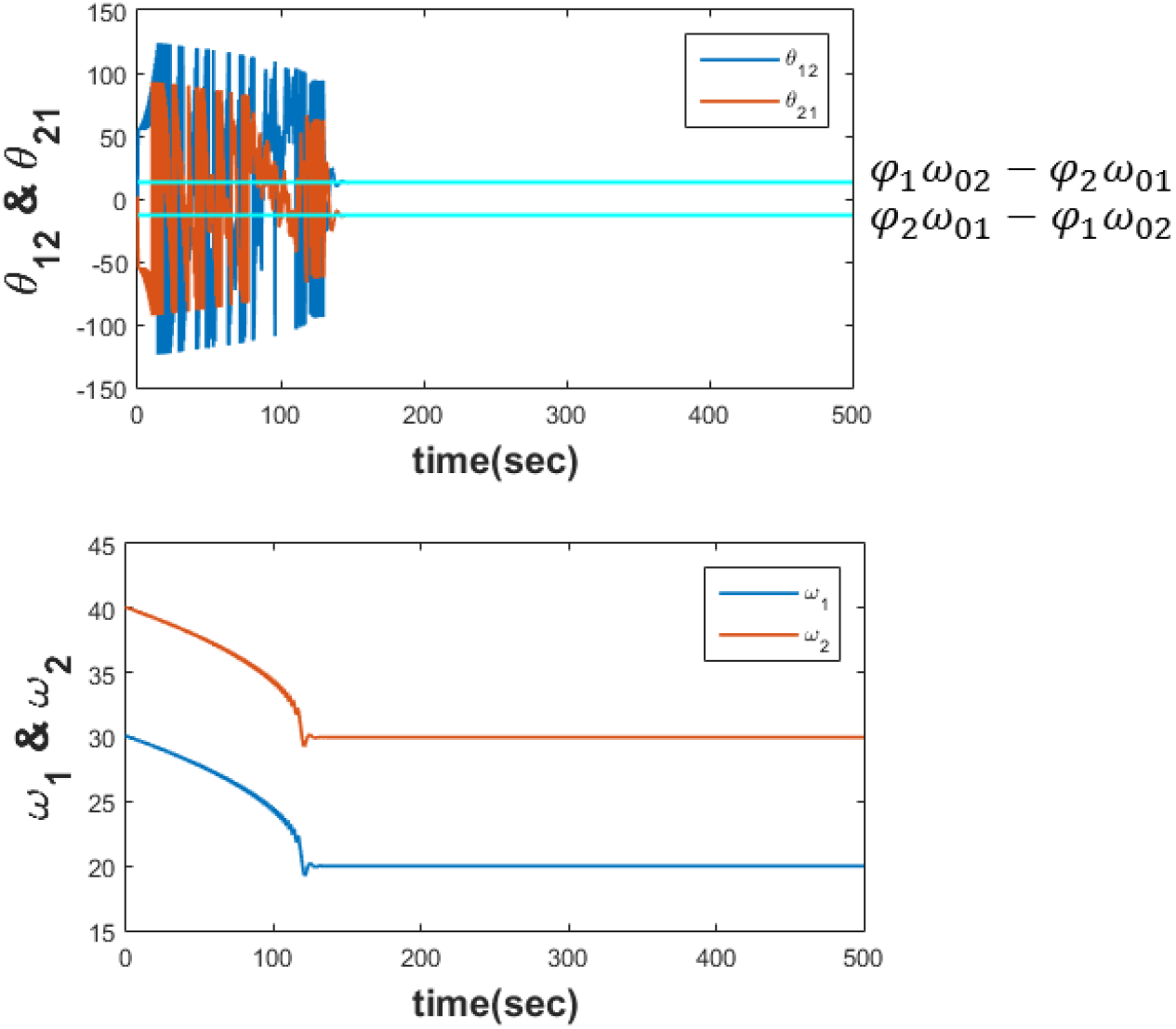
Equations-19a to 19c are simulated for the following set of parameters: *ω*_01_ = 20,*ω*_02_ = 30,*ω*_1_(0) = 30,*ω*_2_(0) = 40,*A*_12_ = *A*_21_ = 0.0001,*θ*_12_(0) = –1.7884,*θ*_21_(0) = 1.7884,*τ_W_* = 10^3^, *ε* = 0.9 and 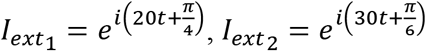. It can be verified that *θ_ij_* learns *φ_j_ω*_0*i*_ – *φ_i_ω*_0*j*_ after the *ω_i_*’s learns the corresponding *ω*_0*i*_’s.

### 2.11 A network for reconstructing a signal by a Fourier-like decomposition

In the previous sections, we have described a network of oscillators in which the natural frequencies and lateral connections can be trained. We now add a feature to the network of Section-2.10 and make it learn an unknown signal by performing a Fourier-like decomposition.

To this end, we construct a reservoir of Hopf oscillators (fig-14). It consists of a network of Hopf oscillators with trainable lateral connections (equation-15) and trainable natural frequencies (equation-20b). In addition, there exists a linear weight stage between the oscillators and the output layer. The natural frequencies of the oscillators adapt themselves to the nearest significant component in the input signal. The lateral connections involving power coupling, encode the (normalized) phase relationships among the oscillators. The output weights represent the amplitudes of the oscillatory components corresponding to the oscillators.

**Fig-14:**
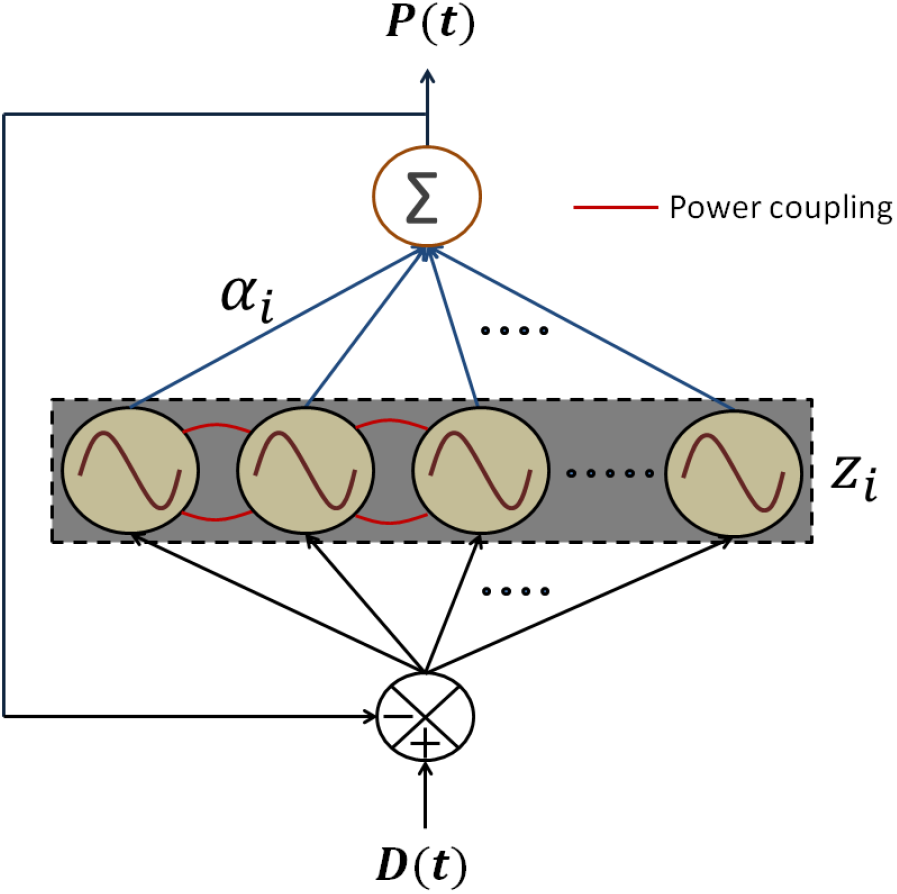
Schematic of the proposed network (equation-20) of a reservoir of N number of adaptive Hopf oscillators similar to the network proposed by (Righetti et al., 2005) with additional asymmetric power coupling connections between the oscillators, each of which is driven by the same error signal *e*(*t*) between the *D*(*t*), teaching time-series signal and *P*(*t*), the linear summation of the oscillations of each of the oscillators in the reservoir through the real feedforward connection weights.

To reconstruct the teaching time-series signal at the output summation node, the error signal that drives each of the oscillators drives the initial phase offset to the desired initial phase offset associated with the corresponding frequency present in the teaching time-series signal. The activation and learning dynamics of the network are given below.

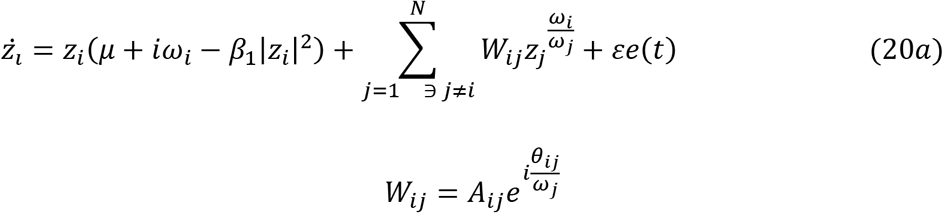

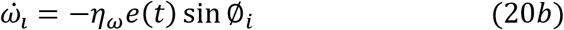

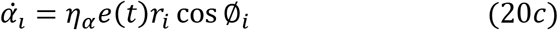

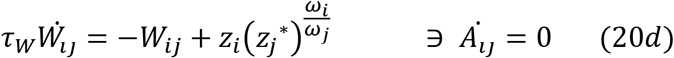

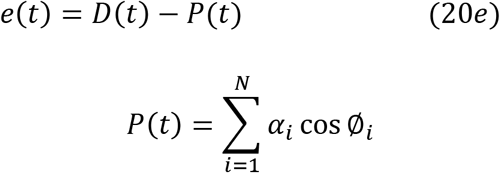

Let *D*(*t*) be the teaching time-series signal with a finite number of frequencies 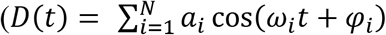 where *φ_i_* be the initial phase offset associated with *i^th^* frequency component), 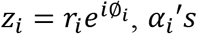 are the real feed-forward weights from *i^th^* oscillator to the output summation node, *τ_W_*, *η_ω_* and *η_α_* are the time constant of the learning dynamics for power coupling coefficient, the learning rate of the natural frequency of the oscillators and the real feed-forward weights from oscillators to the output summation node respectively, *P*(*t*) is the reconstructed signal at the output summation node. The numerical simulation of the proposed network (fig-14 describes the schematic) is elaborated in the following figures (figure-15).

**Fig-15:**
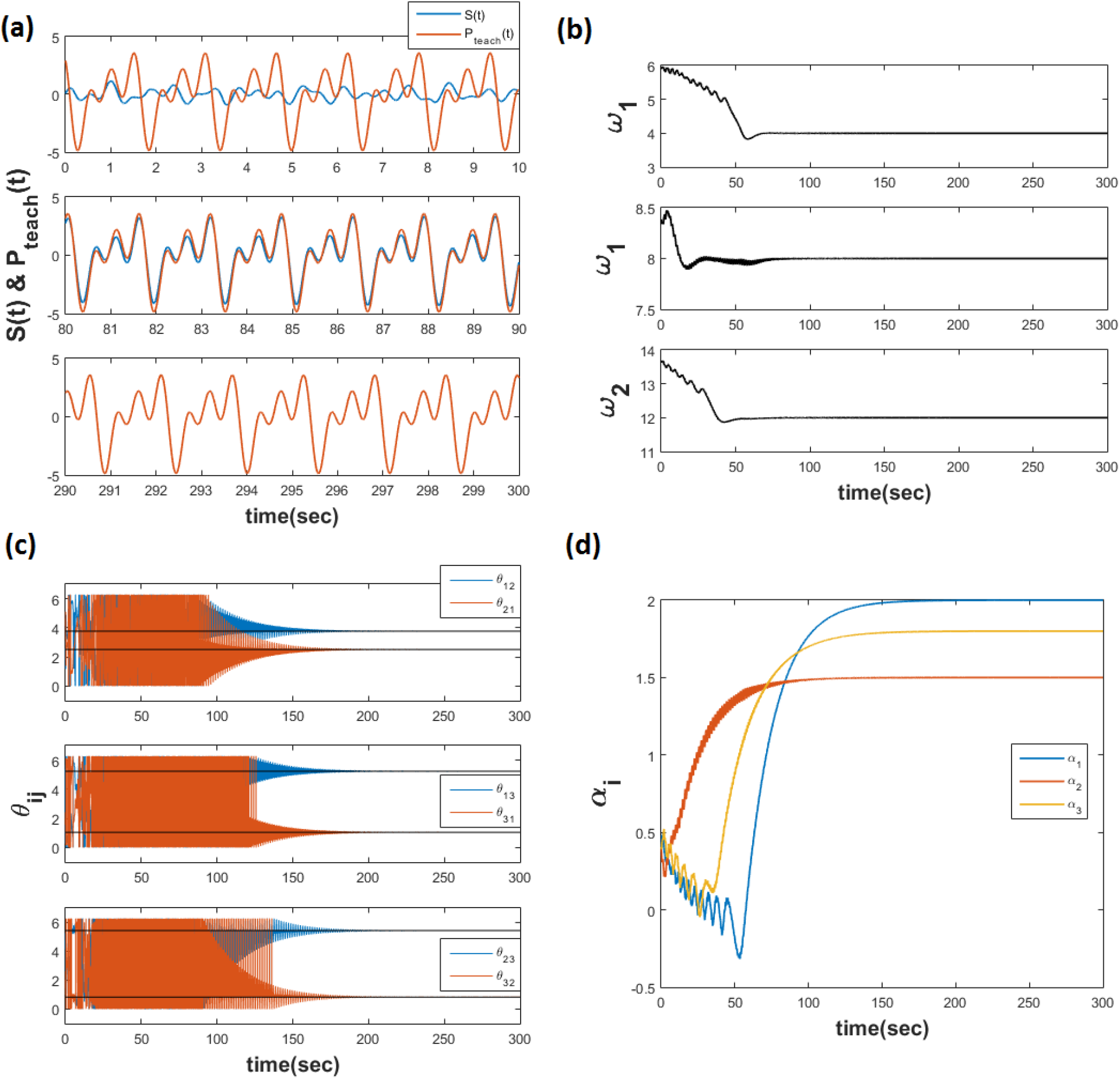
Equations-20a to 20e is simulated with the following parameters 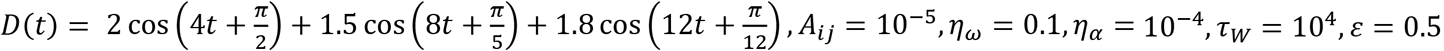, *N* = 3, *dt* = 0.001 *sec*. (a) The network learned signal at the output summation node *P*(*t*) and the *D*(*t*) signal at various 10 secs of time interval. (b) The natural frequencies of the three oscillators learn the three frequency components present in the teaching signal. (c) *ω_j_* times the angles of the complex power coupling weights (*θ_ij_*) learn *φ_i_ω_j_* – *φ_j_ω_i_*. (d) The real feed-forward weights *α_i_*′*s* learn *α_i_*′*s*.

We propose a similar network for Fourier decomposition of complex time series signal with a finite number of frequencies (assuming 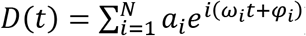), which is an extension of the network proposed in section-2.10. The proposed network is comprised of a reservoir of complex adaptive Hopf oscillators coupled through trainable power coupling weights, and driven by distinct complex sinusoidal inputs with arbitrarily different frequencies. The network is capable of learning the frequencies and encode the normalized phase relationship among the oscillatory components of the input signal. It is driven by the error signal between the complex teaching signal and the linear summation of complex activations of the Hopf oscillators. The dynamics of the network is described in the following equations.

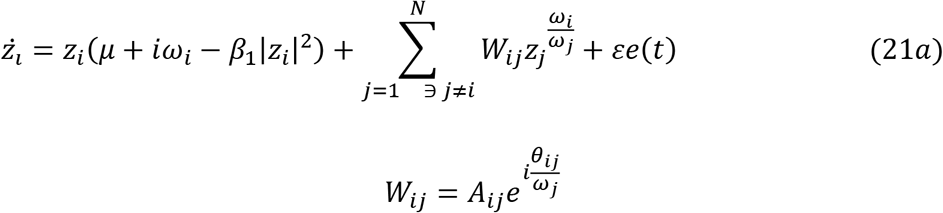

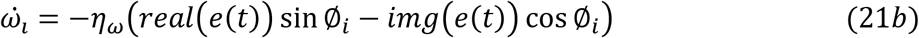

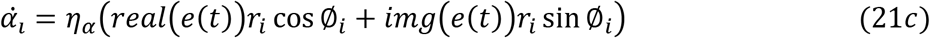

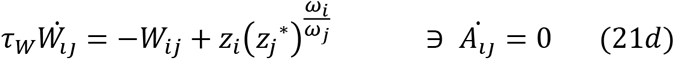

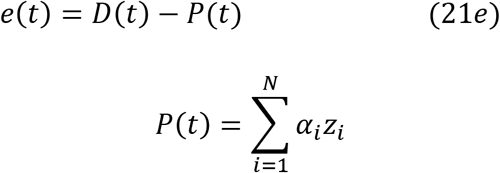

The schematic of the network is identical to the network described in figure-14, and the simulated result for a *D*(*t*), which contains three frequency components is as follows (fig-16).

**Fig-16:**
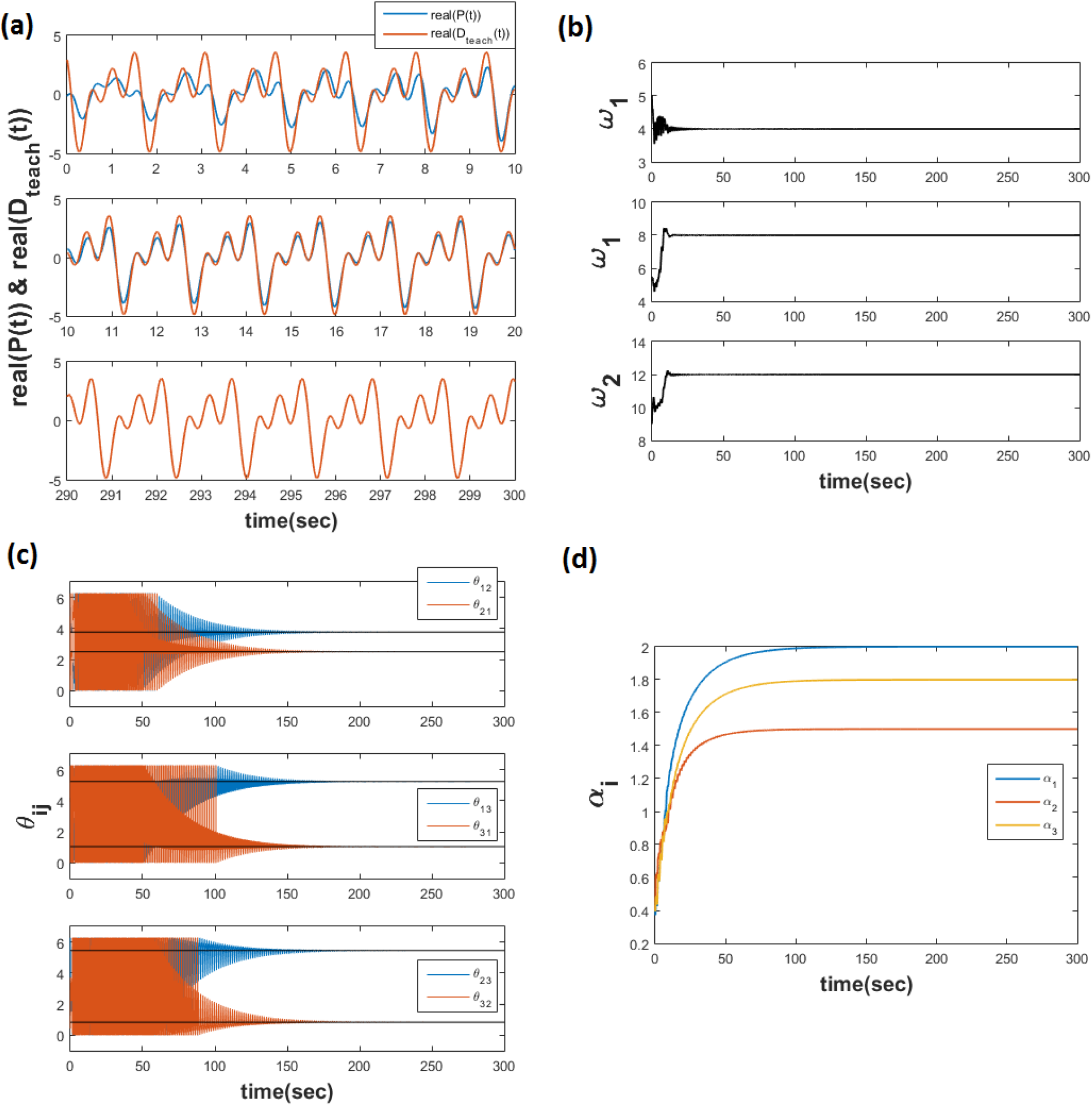
Equations-21a to 21e are simulated for a teaching signal constituting three frequency components with the parameters 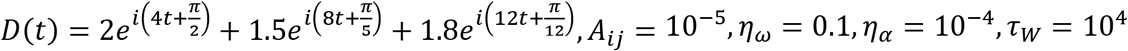, *ε* = 0.5, *N* = 3, *dt* = 0.001 *sec*. (a) The real part of the network learned signal at the output summation node *P*(*t*) and the *D*(*t*) signal at various 10 secs of time interval. (b) The natural frequencies of the three oscillators learn the three frequency components present in the teaching signal. (c) *ω_j_* times the angles of the complex power coupling weights (*θ_ij_*) learn *φ_i_ω_j_* – *φ_j_ω_i_*. (d) The real feed-forward weights *α_i_’s* learn *α_i_’s*.

### 2.12 A generative network which is capable of modeling EEG signals

The proposed network is capable of modeling an arbitrary number of signals with an overlapping frequency spectrum. It is trained in two phases. In the 1^st^ phase, a network exactly similar to the one in the previous section (section-2.11) is used to encode the constituting frequency components of one of the input signals. During this phase, the natural frequencies (*ω_i_*’s), the real feed-forward weights (*α_i_’s*) as well as the power coupling weights 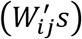 of a network of *N* Hopf oscillators are trained using the same learning rule described in the previous section (equation-20). One key difference compared to the previous section is that the teaching signal in the present scenario is aperiodic (frequency spectrum is continuous) and has a finite duration. To overcome this issue, the limited duration teaching EEG signal is presented repeatedly to the network over multiple epochs. This helps the network to learn some sort of Fourier decomposition of the teaching signal.

In the second phase of training the trained reservoir of oscillators tries to reconstruct *M* number of signals at the corresponding *M* output nodes assuming these *M* signals are the outcome of the same underlying process (producing signals with frequencies confined to a certain frequency band) by training the complex feed-forward weights connecting *N* oscillators of the reservoir to the *M* output nodes. The schematic of the network architecture (fig-17) and the corresponding learning rules are described below.

**Fig-17:**
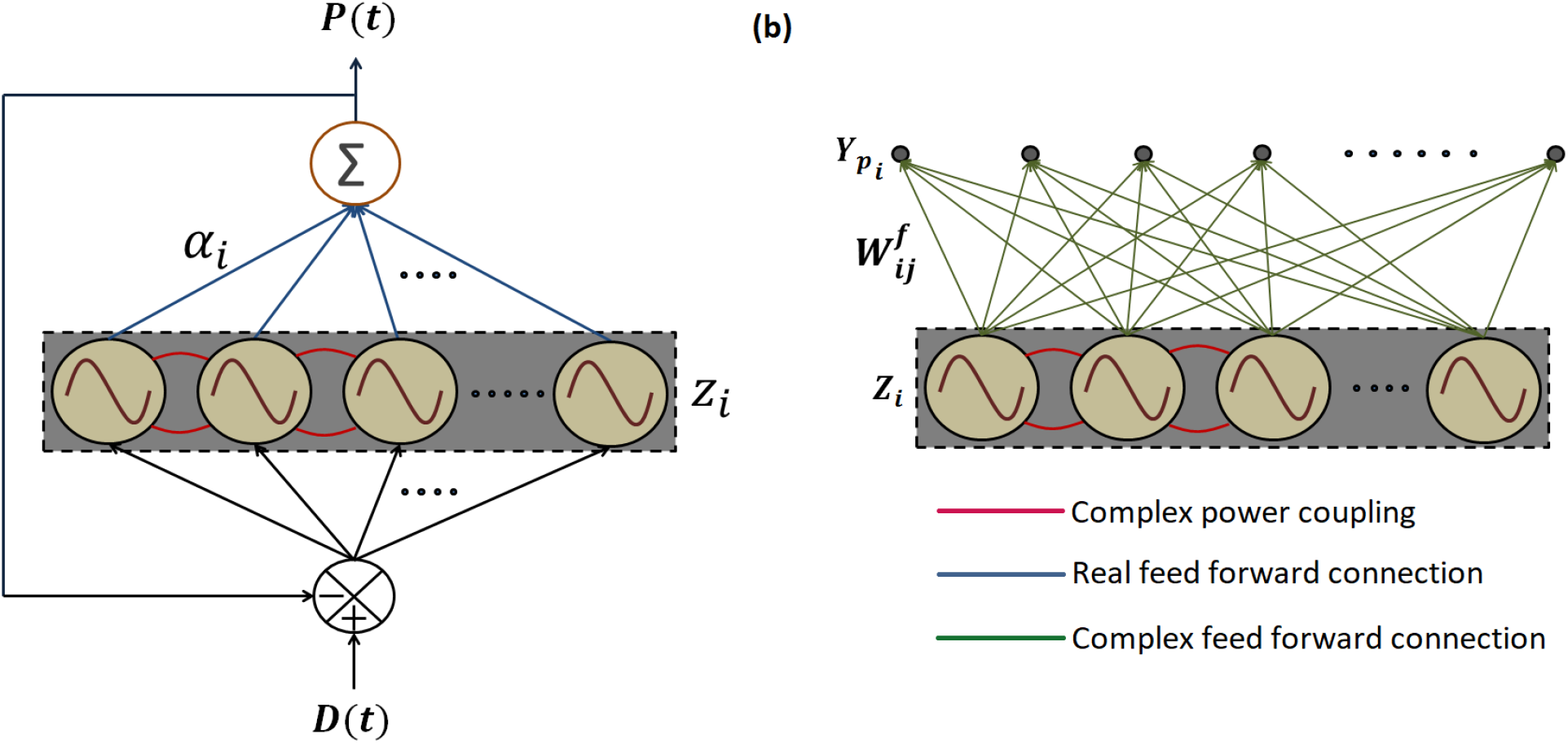
The schematic of the network where (a) elaborates the network architecture used in the first phase of learning, an identical network as used in the previous section elaborated in equation-22. (b) The network architecture of 2^nd^ phase of training with the same reservoir of oscillators with tuned natural frequencies and trained power coupling weights in the 1^st^ phase of training. *Y_p_i__* is the predicted output signal at the *i^th^* output node. Complex feed-forward weights 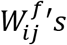 connecting *j^th^* oscillator to the *i^th^* output node, are trained in batch mode according to equation-23.

#### First phase of learning

During the first phase of learning, an identical network with identical learning dynamics as described by equation-20 is used. The only fundamental difference between the previous and present scenario is that *D*(*t*) is a limited duration aperiodic or quasiperiodic signal compared to the previous case where *D*(*t*) was an infinite duration periodic or quasiperiodic signal. i.e., there can be an infinite number of frequency components present in *D*(*t*) as the frequency spectrum of an aperiodic signal is continuous (refer to figure-20). So, the Fourier decomposition using the proposed network can be accomplished by discretely sampling the continuous frequency spectrum.

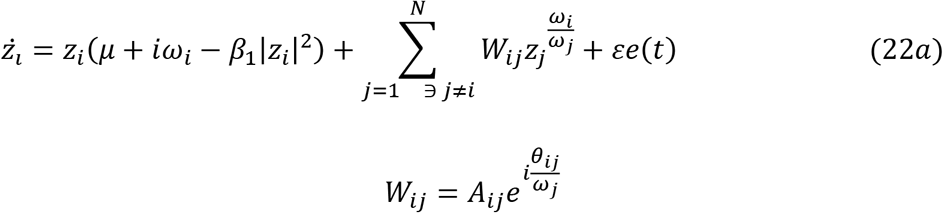

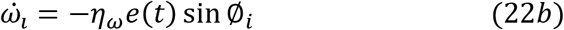

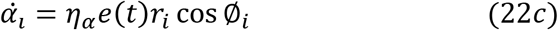

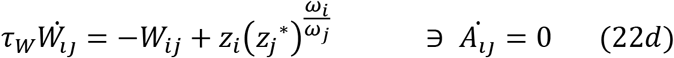

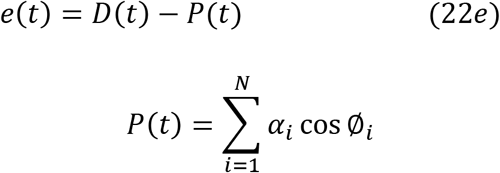

#### Second phase of learning

In this phase, the supervised batch mode of learning is used to train the complex feed-forward weights, *W^f^*, defined as follows. Here *K_ij_* and *ζ_ij_* are respectively the magnitude and angle of the complex feed-forward weights. The derivation for the batch update rule for *k_ij_* and *ζ_ij_* as described in equation-23 is given in the appendix-a5.

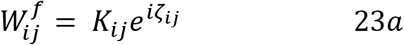

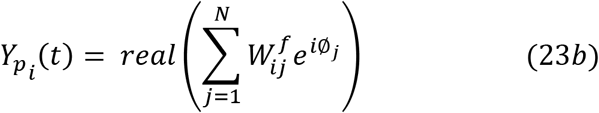

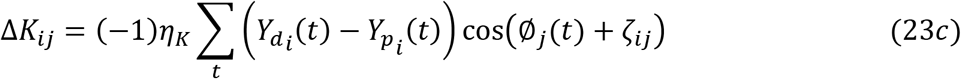

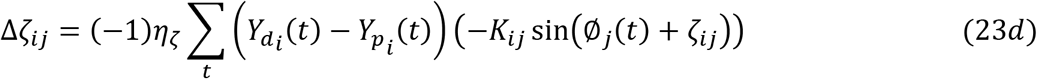

#### Simulation results

At first, the proposed network is simulated to model an arbitrary number of quasiperiodic signals with identical frequency components. The desired signal (*D*(*t*)) consisting of three frequency components (provided 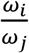 is not an integer) with a duration of 20 secs is used for learning. The network output *P*(*t*) signal (after the first phase of learning) and the respective frequency spectrum is plotted in figure-18. After the second phase of training, the desired and the network reconstructed signals are plotted in figure-19, and the training parameters are described. One crucial condition has to be met in order to achieve an accurate reconstruction: All the oscillators in the reservoir has to be initialized at *z_i_*(0) = 1.

**Fig-18:**
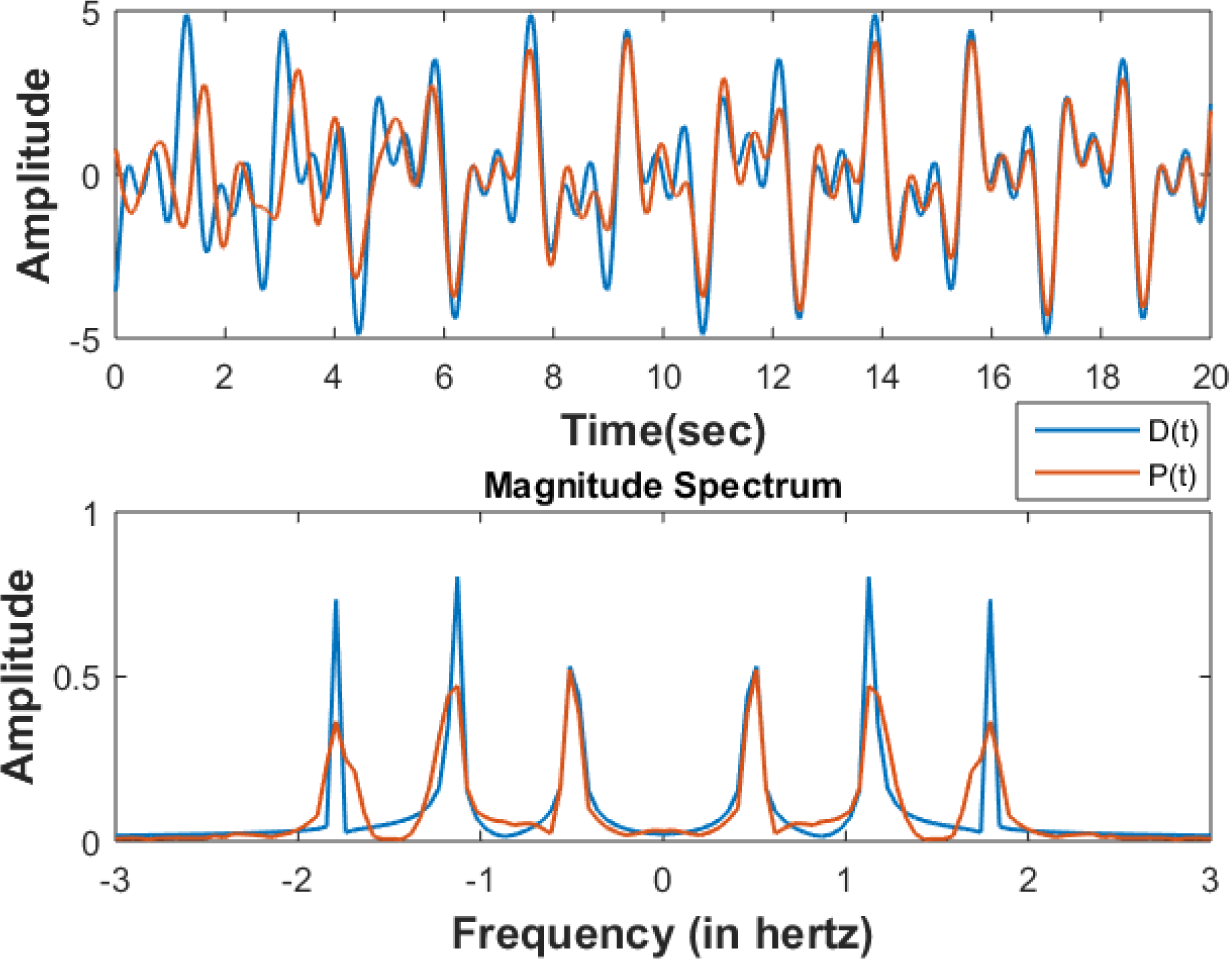
In the first phase of training the oscillatory reservoir network with three oscillators is trained using the following teaching signal and learning parameters: 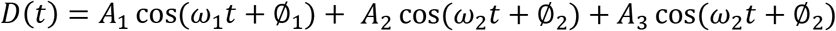, *A_ij_* = 10^-1^, *η_ω_* = 0.1,*η_α_* = 10^-4^, *τ_W_* = 10^4^, *ε* = 0.5, *N* = 3, *dt* = 0.001 *sec, n_epoch_* = 15. The network produced and the original teaching signal and their magnitude spectrum respectively after the last epoch (15) of training.

**Fig-19:**
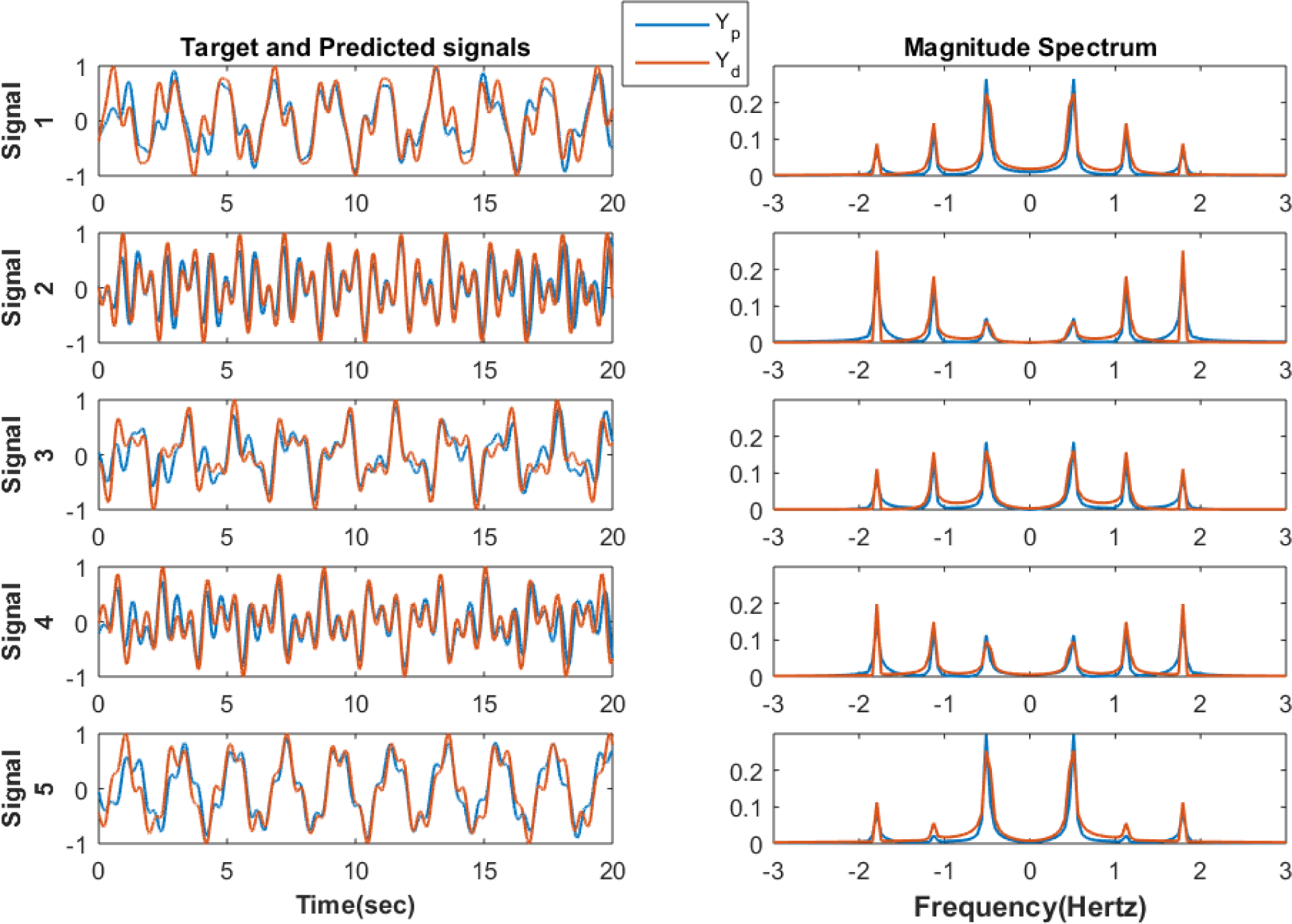
After the second phase of training, the network is able to reconstruct the *M* = 5 output signals (*Y_d_i__* for *i* = 1 to *M*) with identical frequency components but randomly chosen amplitude (*A_i_*) and initial phase offset 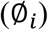 with high accuracy. Learning parameters: *η_κ_* = 3 × 10^-5^, *η_ζ_* = 10^-6^, no. of epochs for the batch mode of learning = 10000.

**Fig-20:**
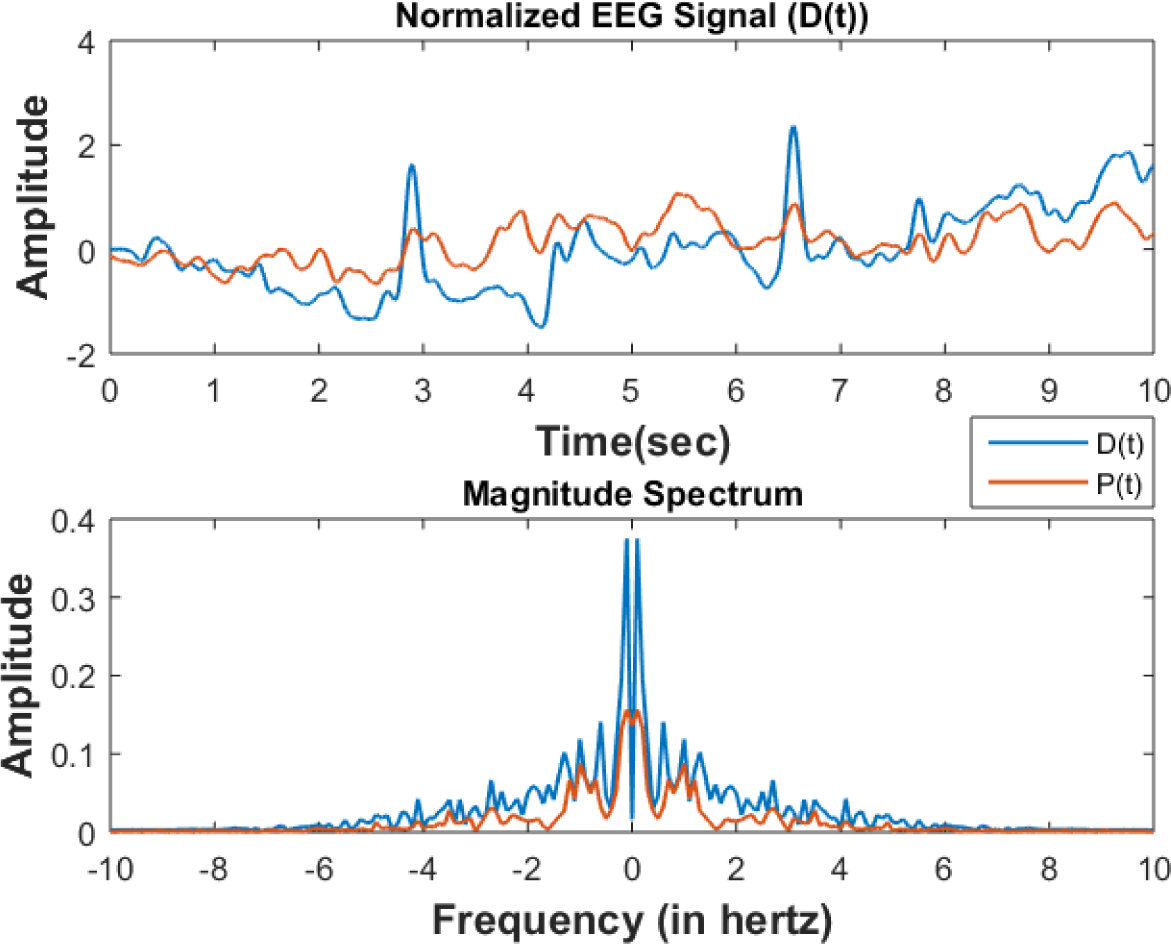
The reconstructed signal by the network of 100 oscillators after 30 epochs, the teaching signal, and the respective frequency spectrum in the first phase of training. In this phase of training, the *D*(*t*) signal is an EEG signal of duration 10 secs, and the exactly similar EEG signal is presented for consecutive 30 epochs. The learning parameters are: *A_ij_* = 10^-1^ *for* |*j* – *i*| ≤ 4 *and A_ij_* = 0 *for* |*j – i*| > 4,*η_ω_* = 0.1, *η_α_* = 10^-4^, *τ_W_* = 10^4^, *ε* = 0.5, *N* = 100, *dt* = 0.001 *sec,n_epoch_* = 30.

The network performance is tested on low pass filtered (cut off frequency of 5 Hertz) EEG data collected from a human subject while performing mind wandering task (Grandchamp et al., 2014). In the first phase, the network encodes *N* number of frequency components of an EEG signal collected from one channel of EEG recording during the mentioned experiment. There is a constraint imposed on the magnitude of the lateral power coupling weights as shown in figure-22. This constraint allows only oscillators with nearby frequency components to interact with each other as initially, the natural frequencies of these oscillators are sorted at an increasing order after sampling from a uniform probability distribution ranging between 0 to 5 Hertz.

**Fig-21:**
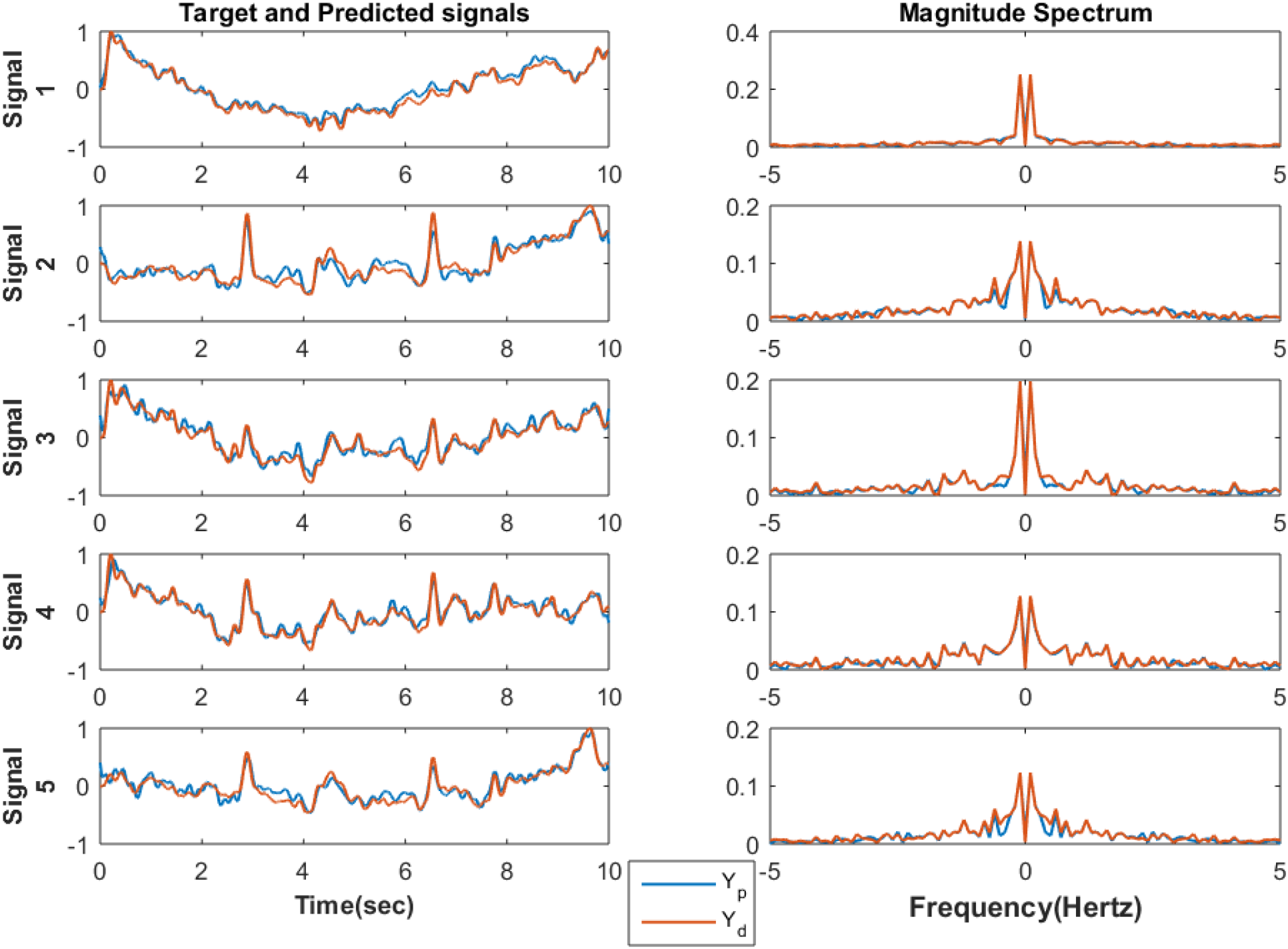
After the second phase of training, the network reconstructed and the original EEG signals collected from 5 channels during the same experiment from which *D*(*t*) for the first phase of training was collected and the corresponding frequency spectrum. Learning parameters: *η_K_* = 3 × 10^-5^,*η_ζ_* = 10^-6^, no. of epochs for the batch mode of learning = 1000.

**Fig-22:**
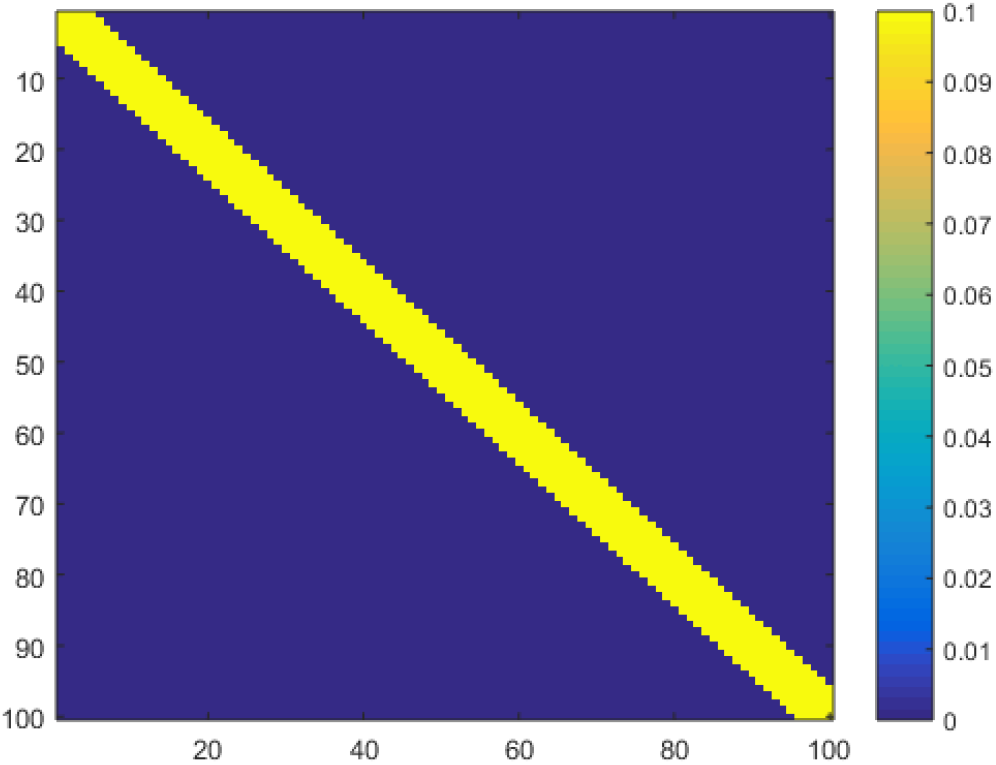
The figure is depicting *the A_ij_* matrix from which it can be seen that a given oscillator is only power coupled with nearby oscillators, which ensures a given oscillator to interact with oscillators having close natural frequency. *A_ij_* = 10^-1^ *for* |*j* – *i*| ≤ 4 *and A_ij_* = 0 *for* |*j* – *i*| >4

In the second phase, the network tries to reconstruct the EEG signals collected simultaneously from the other channels during the same experiment. A set of EEG signals from 5 other channels are reconstructed by the network. The reconstruction accuracy of the modeled EEG signals depends on the number of oscillators in the oscillatory layer and the number of desired signals to be reconstructed. Using complex weights for the feed-forward connections is a key factor as it helps to learn the amplitude, as well as phase of the Fourier decomposition of the EEG signal to be reconstructed, provided the frequency components are already learnt in the encoding phase.

## 3. Discussion

The unique contribution of the proposed network of complex neural oscillators is the notion of power coupling by which it becomes possible to achieve a stable normalized phase relationship between two oscillators with arbitrary natural frequencies. With such a feature as the backbone, it is possible to construct a network of oscillators that can learn to reconstruct multiple time series signals. Another positive feature of the proposed model is the biological feasibility of the learning mechanisms. The lateral connections in the oscillatory reservoir are trained by a Hebb-like rule, while the forward connections are trained by a modified delta rule adapted to the complex domain.

Network models of complex-valued oscillators have been described before. For example, the result that through a complex coupling, it is possible to achieve an arbitrary phase difference between two oscillators with equal natural frequencies was described in (Hoppensteadt and Izhikevich, 2000). However, Hoppensteadt and Izhikevich did not extend the result to the case of interacting oscillators with very different natural frequencies, as we have done using power coupling. A network of complex-valued oscillators that can store patterns as oscillatory states was described in (Chakravarthy and Ghosh, 1996). The model also proposed a complex form of Hebb’s rule, similar to the one used in the current study. However, the model of (Chakravarthy and Ghosh, 1996) was limited in its capability by the fact that all the oscillators in the model have a common frequency. A representation of synaptic strength using a complex weight, instead of the usual real weight, has an added advantage in representing the temporal relationships underlying neural dynamics. We have recently shown (Chakravarthy, 2020) that the imaginary part of the complex weight captures the temporal asymmetry between the activities of pre- and post-synaptic neurons in a manner akin to the weight kernel of Spike Time Dependent Plasticity (STDP) mechanism (Bi and Poo, 1998).

It may be said that the network of Section 2.11 achieves a discrete approximation of the continuous spectra of the signals that are reconstructed by the network. The greater the number of oscillators in the network, the tighter is the approximation. In this approximation, the trainable real feed-forward weights (*α_i_*) of the network learn the amplitude spectrum of *D*(*t*). The *θ_ij_* of the power coupling weight learn normalized phase differences, *φ_i_ω_j_* – *φ_j_ω_i_*.

It has previously been shown in (Righetti et al., 2005) that, when a network of supercritical Hopf oscillators driven by the reconstruction error signal, individual oscillators are tuned to the nearest frequency components in the target time series. In the network (net-1: schematic is described in figure-2, and the network dynamics and the learning dynamics is given in equation-4 to 8 of Righetti et al., 2005) one shortcoming is that there is no way to recreate the teaching timeseries signal (*P_teach_*(*t*)) back without the error feedback loop as with the learnt *α_i_*’s and *ω_i_*’s, when the network is reset (i.e., *r_i_*’s and 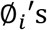 are reset) there is no external forcing to drive the initial phase offset of each oscillator to the desired phase offsets 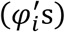. To overcome this issue, the authors have proposed a phase signal of frequency same as *i^th^* oscillator but same phase as the oscillator with the lowest frequency as a signal communicated by the oscillator with the lowest frequency to the *i^th^* oscillator, which makes those two oscillators maintain a certain normalized phase difference 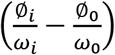. However, to learn/store the phase relationship between the lowest frequency component and the *i^th^* frequency component in the teaching signal a “phase variable” considered as a third variable of the oscillator dynamics has been defined which learns the difference between phase signal from the oscillator with the lowest frequency and its own phase. The main issue with this network is that the network can only learn Fourier decomposition (frequency, magnitude and relative initial phase offset among the constituting frequency components) of a periodic signal which has frequency components of the form *ω_i_* = *nω*_0_, *ω*_0_ being the fundamental frequency of the periodic signal as *n∈N* as the phase signal *R_i_* from the 0^*th*^ oscillator is interacting with the *x* dynamics of the 0^*th*^ oscillator through real coupling constant. Furthermore, there is an ambiguity in defining the lowest frequency component or fundamental frequency of the periodic teaching signal as the dynamics do not itself identify the oscillator, which learns the lowest frequency component in the teaching signal.

To overcome these issues, the proposed network as described in section-2.11 is a similar network where the reservoir of complex supercritical Hopf oscillators is coupled through power coupling to preserve the phase relationship among the constituting frequency components in the teaching signal. When second network architecture proposed by Ijspeert et al. (schematic of which is described in figure-3 and the dynamics is given in equation-13 to 20 of Righetti et al., 2005) tries to learn Fourier decomposition of a teaching signal the natural frequencies of the oscillators and the real feed-forward weights learns the magnitude spectrum of the teaching signal. During learning the error signal, *F*(*t*) drives the initial phase 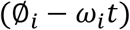 of each of the oscillators to the desired initial phase (*φ_i_*) can be retrieved from the phase spectrum of the teaching signal. Thus the same network deliberated by equation-20 with the oscillators coupled through power coupling with fixed *A_ij_*(≪ 1) ensures the angle of power coupling coefficient (*θ_ij_*) to learn *φ_i_ω_j_* – *φ_j_ω_i_*.

One issue while retrieving the stored oscillatory pattern is the dependence on the initial state of the oscillators to achieve the desired solution (see section-2.8). We observed that when 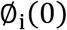’s are sampled from space w subspace of W = {*x* ∈ *R^N^*|0 ≤ *x* < 2*π*}, the network attains the desired solution at a steady-state. However, there are other plausible solutions or spurious states the network can acquire. The desired solution of *σ_ij_*\*(= 0) and the spurious solutions are independent of the network parameters like *A_ij_* (for *A_ij_* = *A*) and *θ_ij_* but has a dependence on the natural frequency of the oscillators. However, the solutions of *ψ_ij_* are dependent on both network parameters (*θ_ij_* and *ω_i_*, assuming *A_ij_* = *A*). The dependency of the boundary of space ω or the boundary of the other subspaces (initializing 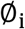 by sampling which leads the system to spurious states) on the natural frequency of the oscillators is yet to be understood.

There is a vast literature pertaining to complex-valued neural network models, which are produced by extending their real-valued counterparts to the complex domain. Complex feedforward networks trained by the complex backpropagation algorithm, complex Hopfield network, complex self-organizing maps are examples of such models. Complex formalism plays a key role in linear systems theory in which both systems and signals can be represented as complex functions. As a special case of this, in electrical circuit theory, oscillations are modeled as complex numbers called phasors, that represent the phase of the oscillations. Therefore, there is a longstanding association between complex numbers and oscillations. Nevertheless, complex-valued neural network literature did not seem to have exploited this association. For the most part, complex-valued neural networks operate like 2n dimensional versions of their n-dimensional real-valued counterparts. The proposed model attempts to take an important step in filling this lacuna. By describing oscillations in the complex plane, and invoking the power coupling principle, it is able to elegantly overcome some of the difficulties involved in harnessing the dynamics of multi-oscillator networks with real coupling.

In the present study, we use the oscillatory neural network model to reconstruct six EEG time series simultaneously. However, the proposed learning mechanisms and network architecture can be easily scaled up. It is possible to model a much larger number of EEG channels (say, 128 or 512) simply by increasing the size of the oscillatory reservoir and the number of output neurons. In the current study, the connections among the oscillators are constrained to ensure that only oscillators with nearby natural frequencies are connected. In a large-scale model, it is conceivable to associate the individual neural oscillators with anatomical locations in the brain, and constrain their coupling based on structural connectivity information from structural imaging tools like Diffusion Tensor Imaging (DTI) (Damoiseaux and Greicius, 2009). Through such developments, the present model can evolve into a class large-scale models of brain dynamics similar to The Virtual Brain (TVB) model, with some relative advantages.

The TVB model principally uses neuronal mass models like the two-variable nonlinear oscillatory models including Fitzhugh-Nagumo model, Winson-Cowan model, Wong-Wang model, Brunel-Wang model, Jensen-Rit model, Stefanescu-Jirsa model (Wilson and Cowan, 1972; Wong and Wang, 2006; Brunel and Wang, 2003; Jansen and Rit, 1995; Stefanescu and Jirsa, 2008), which can exhibit excitable dynamics and limit cycle oscillations. In these models, the oscillators are coupled by only one variable, which is analogous to “real-valued” coupling. Furthermore, most of them do not have an explicit natural frequency parameter that is trainable. Therefore, the coupling connections cannot be trained using a biologically feasible learning rule like Hebbian learning. They are trained by global optimization algorithms that adjust the coupling strengths so that the output of the oscillatory network matches the recorded brain dynamics.

In the future, we plan to extend the proposed model to create a whole class of deep oscillatory networks. The network will have an input stage that serves as an encoder that performs a Fourier-like decomposition of the input time-series signals. The network model of Section 2.11 will play the role of this encoder. The hidden layers will consist of oscillators operating at a range of frequencies. The output layer will be a decoder that converts the oscillatory outputs of the last hidden layer into the output time series. Another interesting proposed study is to develop the network model of Section-2.11 into a model of the tonotopic map. Electrophysiological recordings from the bat’s auditory cortex revealed a map of frequencies (a tonotopic map). Kohonen had proposed a model of the tonotopic map using a self-organizing map (Kohonen, 1998). However, this model represents frequency as an explicitly available parameter, without actually modeling the responses of the neurons to oscillatory inputs (tones). We propose that by organizing the oscillators in the reservoir model of Section 2.11 in a 2D array with neighborhood connections, it is possible to produce a biologically more feasible tonotopic map model.

## Acknowledgements

We acknowledge the support from undergraduate student from Mechanical Department of IIT Madras, Asit Tarsode for his insight on the dependency of steady state solutions on the initial conditions. We also acknowledge the constant and much needed support from senior lab mate Vignayanandam R Muddapu, and other lab members.

## Appendix

## A1. A pair of coupled Hopf oscillators with real coupling

The dynamics of a pair of Hopf oscillators represented in equation-2 is given below:

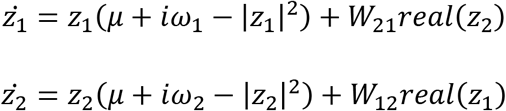

Assuming *ω*_1_ = *ω*_1_ = *ω*, and W_12_ and W_21_ are real and small, the polar coordinate representation is:

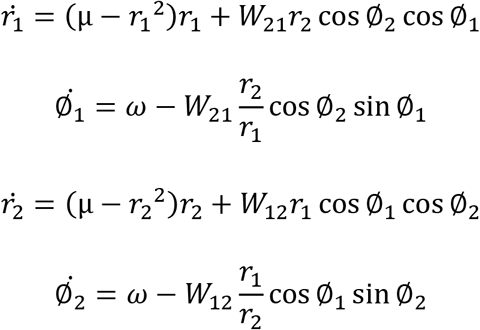

Let the phase difference between the oscillators, 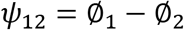,

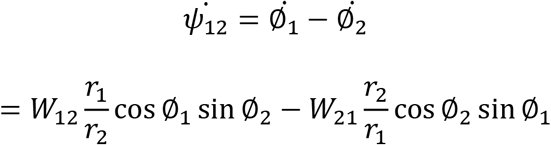

At steady state, 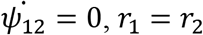,

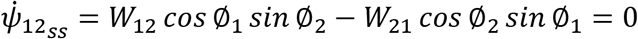

When *W*_12_ = *W*_21_ = *ε*, a real number,

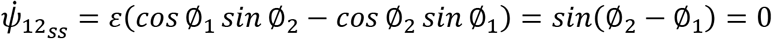

i.e. the solution is: 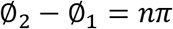, *n* being an integer.

When, *ε* is a positive real number, 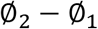 attains 2*nπ* solutions, whereas if *ε* is negative real number 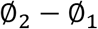 attains (2*n* + 1)*π* solutions.

## A2. Proof for phase-locking using complex coupling coefficient

To ensure two Hopf oscillators with identical natural frequencies be phase-locked at any particular angle independent of the initial condition, the two oscillators need to be coupled using a complex coupling coefficient. With the coupling strategy, as stated in equation-4, it is analytically shown below that the at steady-state two Hopf oscillators will be phase-locked at a particular angle that equals the angle of complex coupling coefficient.

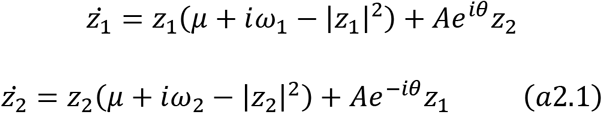

Let 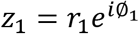 and 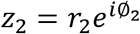 Where *α* = *μ* (> 0)*β*_1_ = −1, *ω*_1_ = *ω*_2_ = *ω* small A;

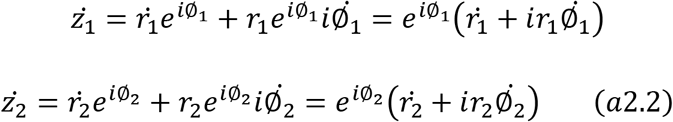

Substituting equation-a2.2 in equation-a2.1,

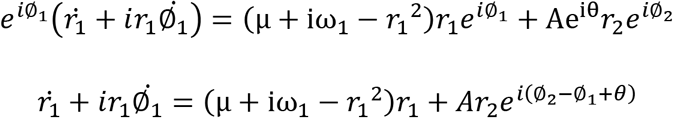

Similarly,

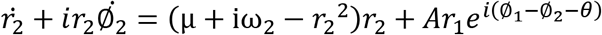

Equating real and the imaginary term independently,

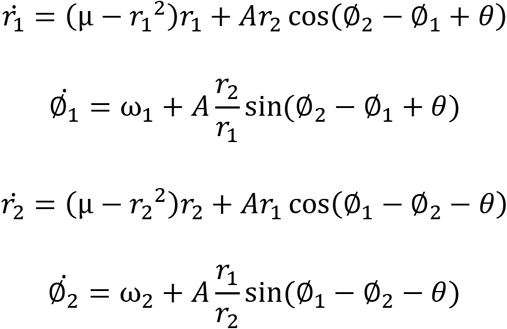

Let *ψ* be defined as 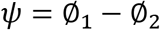

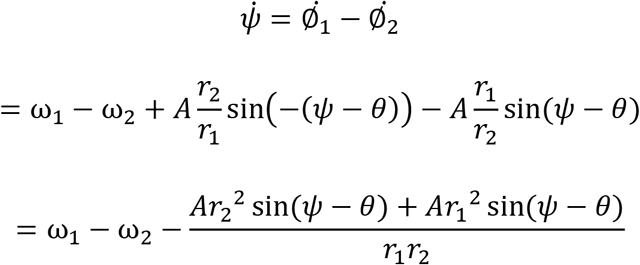

In this scenario *ω*_1_ = *ω*_2_ and at steady state 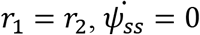,

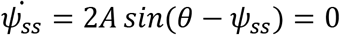

Implying that,

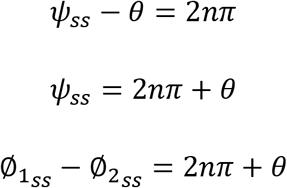

## A3. Two complex Hopf oscillators with power coupling

Consider two supercritical Hopf oscillators with different natural frequencies coupled as described in the following equations. We explore under the steady-state condition, what would be the phase relationship of the two oscillators.

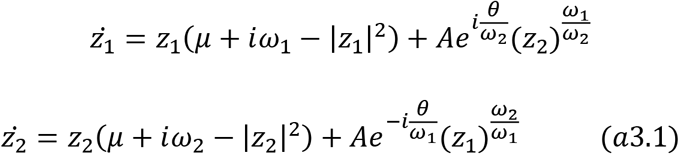

Let 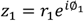 and 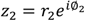, where *α* = *μ* (> 0), *β*_1_ = − 1,

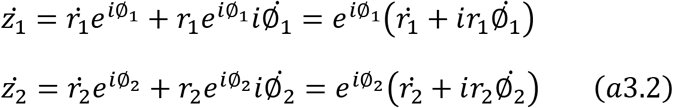

Substituting equation-a3.2 into equation-a3.1,

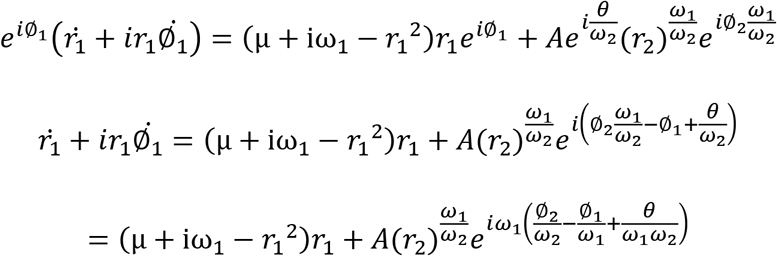

Similarly,

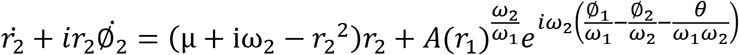

Equating the real and imaginary terms,

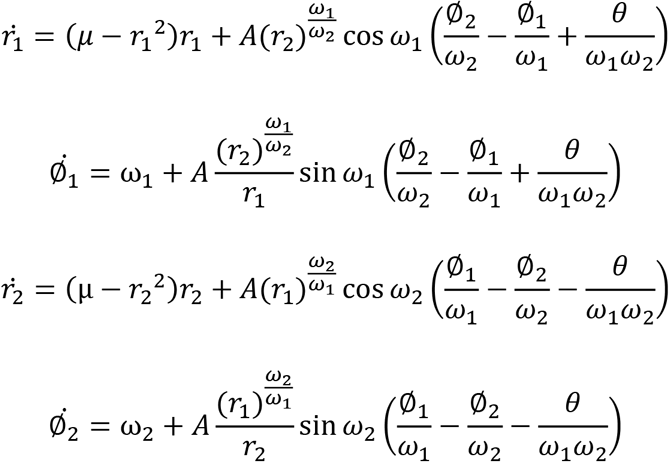

At steady-state, the relationship between two oscillator’s natural frequencies *ω*_1_ and *ω*_2_ and phase 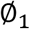 and 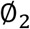 respectively holds the following relation,

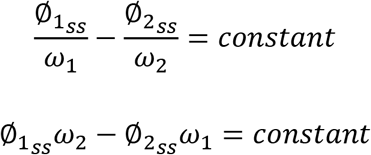

This constant can be any real number. Let’s denote *ψ* as 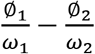, as we know that *ψ* is constant at steady state, hence 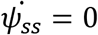

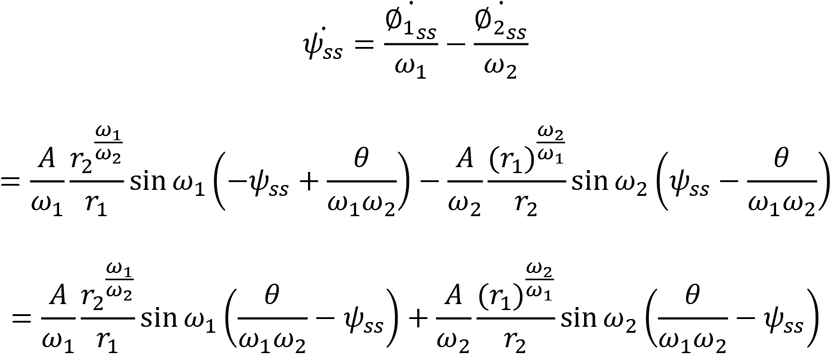

At steady-state *r*_1_ = *r*_2_ as *μ* for both of the oscillators equals 1, the previous equation becomes,

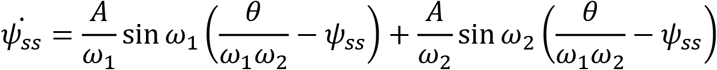

As 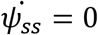, implying that, for a subset (say w) of solutions:

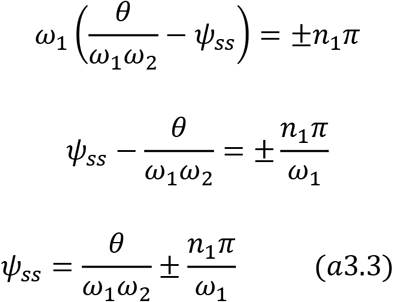

as well as,

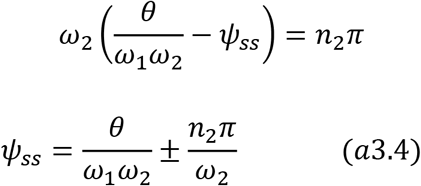

From the above equation-a3.3 and a3.4, when *n*_1_ = *n*_1_ ≠ 0 (other than the desired solution for which *n*_1_ = *n*_1_ = 0), the subset w will only contain desired solution when 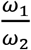 is an irrational number.

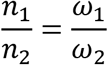

## A4. Derivation of learning rule for power coupling weight

If *N* of supercritical Hopf oscillators is laterally connected with each other through power coupling, the dynamical equations will look like:

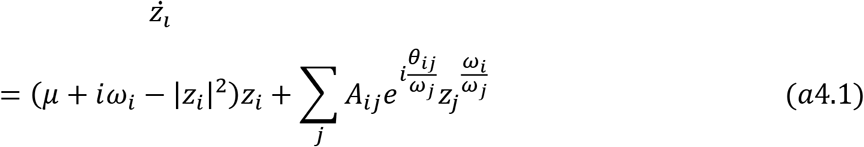

where 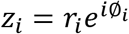

Therefore,

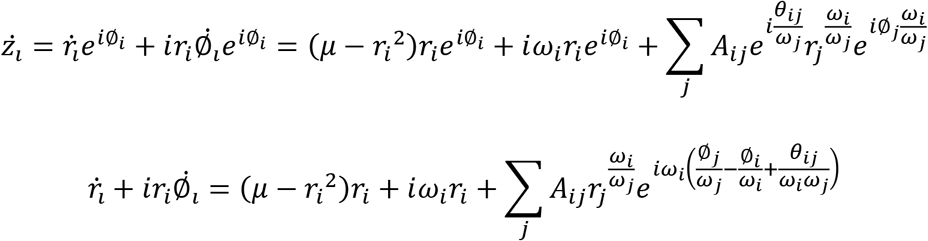

Separating real and imaginary part;

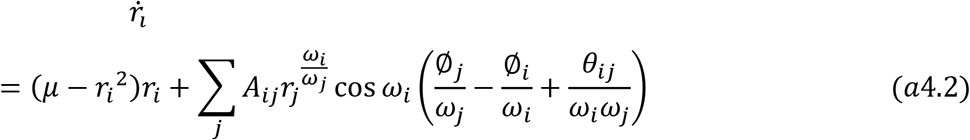

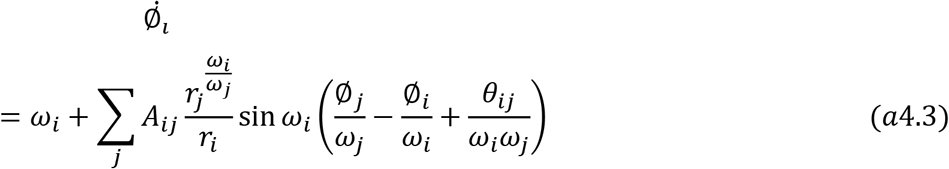

The normalized phase difference between *i^th^* and *j^th^* Hopf oscillator,

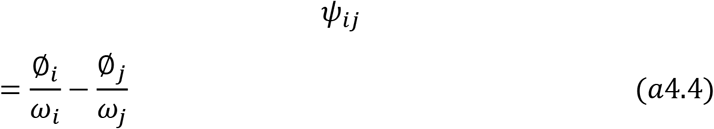

Therefore, from equation-a4.3 and a4.4,

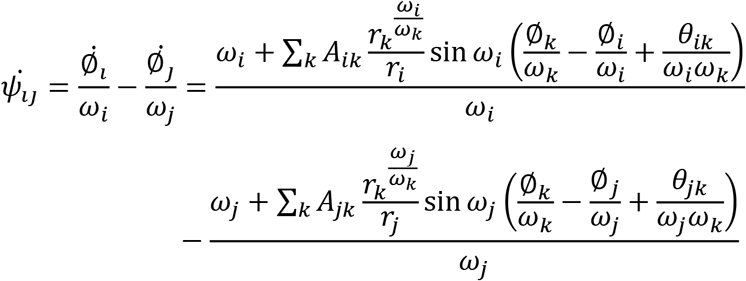

Or,

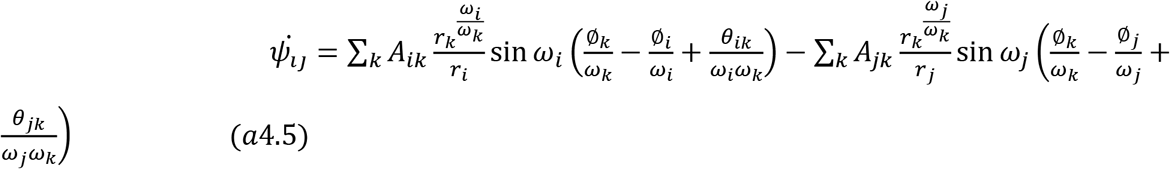

At steady state, 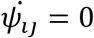
and, 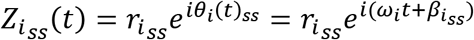
where *β_i_ss__* and *r_i_ss__* are the steady state phase offset and magnitude of the *i^th^* oscillator.

From equation-a4.5,

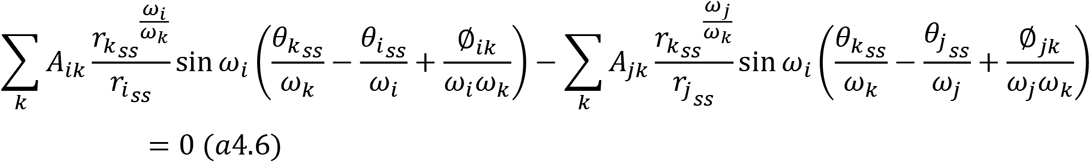

One obvious of equation-a4.6 is,

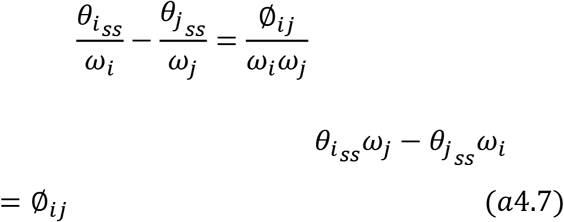

Thus, at the time of learning the Fourier decomposition of a given perturbing input signal, to restore the normalized phase of oscillations of the Hopf oscillators 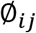, the angle of complex lateral coupling has to be updated according to equation-a4.7, which boils down to the following learning rule of the lateral connection:

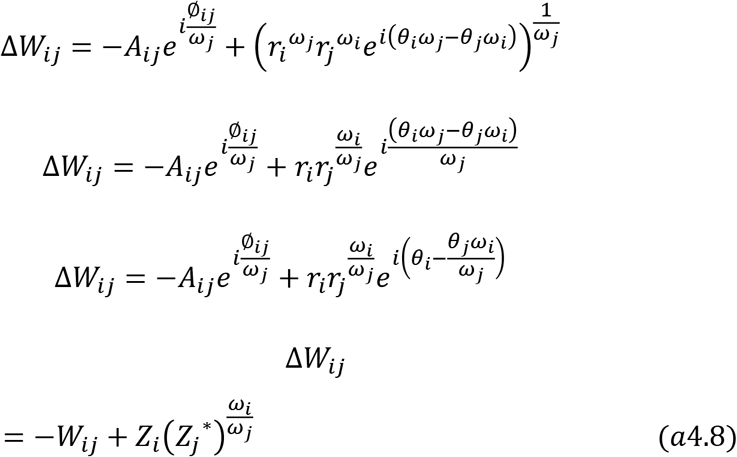

## A5. Derivation of the complex feed-forward weights in the generative network

For the batch mode of learning the squared loss function is as defined;

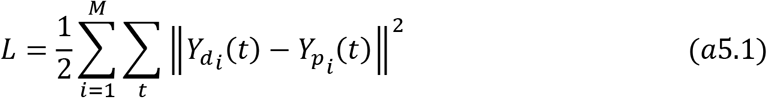

where,

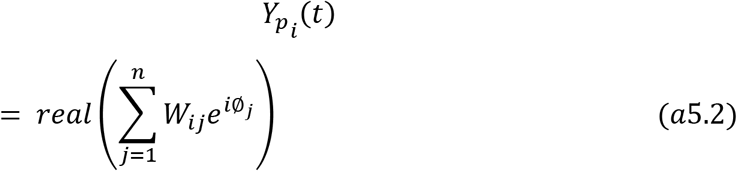

and *W_ij_* is the complex output weight from *j^th^* Kuramoto oscillator to the *i^th^* output node, defined as;

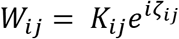

From equation-2,

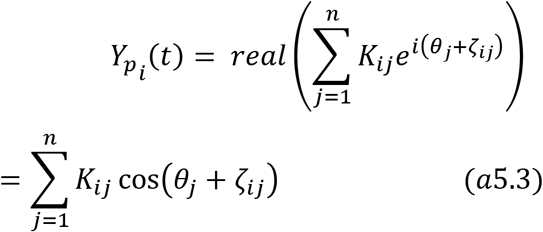

From equation-a5.1 and a5.3 the update rule for *K_ij_* and 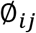 can be derived as;

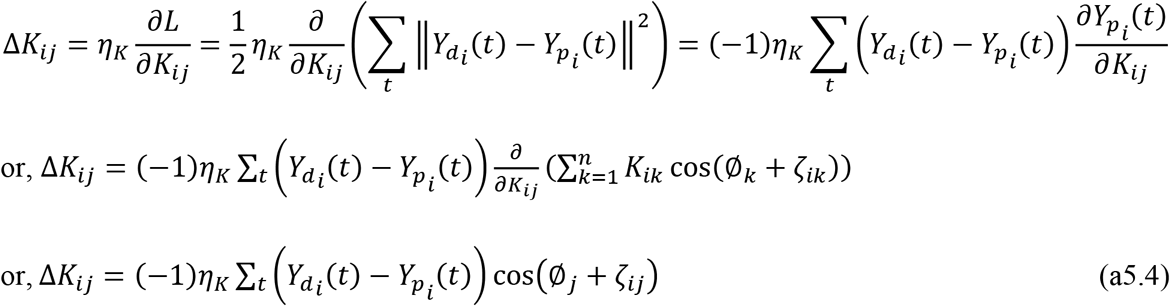

Similarly;

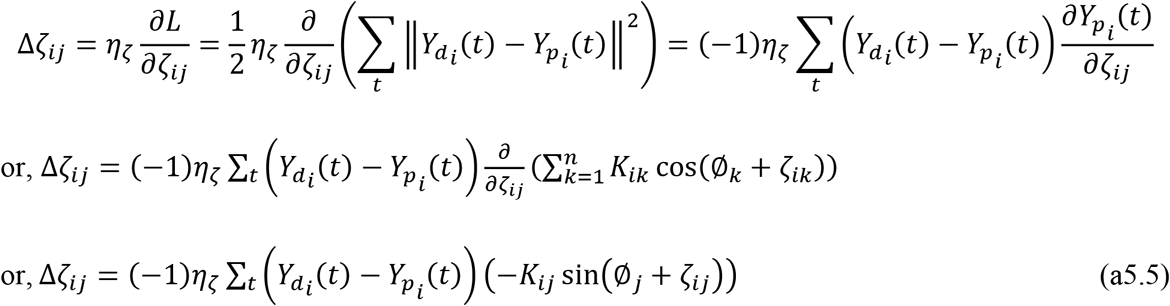

